# Disruption of the epigenetic regulator BAP1 drives chromatin remodeling leading to the emergence of cells with breast cancer stem cell properties and aberrant glycosylation

**DOI:** 10.1101/2024.12.12.628129

**Authors:** Mariana Gomes da Silva Araujo, Aurélie Sallé, Vincent Cahais, Claire Renard, Cyrille Cuenin, Caroline Pires Poubel, Stéphane Keita, Thorsten Mosler, Christine Carreira, Gabrielle Goldman Levy, Lya Parres, Ekaterina Bourova-Flin, Sophie Rousseaux, Saadi Khochbin, Akram Ghantous, Siniša Habazin, Maja Pučić-Baković, Erika Cosset, Gordan Lauc, Zdenko Herceg, Rita Khoueiry

**Affiliations:** International Agency for Research on Cancer, World Health Organization, 69007, Lyon, France; Institute of Biochemistry II, Faculty of Medicine, Goethe University Frankfurt, Frankfurt, Germany; Université Grenoble Alpes, CNRS UMR 5309, INSERM U1209, Institute for Advanced Biosciences, 38700, Grenoble, France; Genos Glycoscience Research Laboratory, 10000, Zagreb, Croatia; Cancer Research Center of Lyon - CRCL, INSERM U1052, CNRS UMR 5286, 69008, Lyon, France; Faculty of Pharmacy and Biochemistry, University of Zagreb, 10000, Zagreb, Croatia

**Keywords:** Breast Cancer, Epigenetics, CRISPR screening, Glycosylation, Breast Cancer Stem Cells

## Abstract

**Background:** Epigenetic regulator genes play critical roles in controlling cell identity and are frequently disrupted in breast cancers, suggesting a key driver role in this disease and its associated phenotypes. However, specific epigenetic drivers (epidrivers) of mammary cell plasticity and their mechanistic contributions to this phenotype are poorly characterized.

**Methods:** To identify potential epidrivers of the emergence of mesenchymal breast cancer stem cell-like phenotypes in non-tumorigenic mammary cells, we employed a CRISPR/Cas9 loss-of-function screening strategy targeting epigenetic regulator genes. This approach was followed by an in-depth validation and characterization of epigenomic, transcriptomic, proteomic and phenotypic changes resulting from the disruption of the putative epidriver gene *BAP1*.

**Results:** Our investigation revealed that loss of the histone deubiquitinase BAP1 impacts cellular processes associated with breast cancer cell plasticity such as epithelial-to-mesenchymal transition (EMT) and actin cytoskeleton organization. In addition, we unveiled that BAP1 loss resulted in an overall less permissive chromatin and downregulated gene expression, impacting programs that control cellular glycosylation and leading to decreased glycan abundance and complexity. BAP1 rescue restored the expression of several deregulated genes in a catalytic activity-dependent manner, suggesting that BAP1-mediated cell identity and glycosylation regulation are largely dependent on its histone deubiquitinase activity.

**Conclusions:** Overall, our results point to BAP1 disruption as a driver of mammary cell plasticity and reveal a novel role of BAP1 as an epigenetic regulator of cellular glycosylation.

## BACKGROUND

International wide-ranging efforts in sequencing cancer genomes and epigenomes have revealed a high frequency of alterations in genes encoding proteins that regulate the epigenome (epigenetic regulator genes, ERGs) across different cancer types ^1,2^. This striking discovery underscored the importance of epigenetic mechanisms in cancer causation and biology. In a recent study, we showed that the disruption of ERGs may be associated with cancer hallmarks in many cancer types, supporting the notion that ERG disruption underpins rampant epigenome alterations in cancers ^3^. However, the timing and stage at which the disruption of ERGs assume a primary role in tumorigenesis are poorly understood.

In a cancer setting, epigenetic deregulations largely contribute to oncogenic reprogramming, enhanced cell plasticity, and maintenance of specific cancer cell phenotypes ^4,5^. Therefore, epigenetic aberrations may enable tumorigenesis through the emergence of cancer stem cells (CSCs) or cancer initiating cells, and underlie tumor promotion and progression by conferring specific properties to cells ^6,7^. In breast cancer (BC), subpopulations of highly plastic cells which can transition into different CSC states have been described. These breast CSCs (BCSCs) contribute to tumor growth, invasion, and resistance to treatment ^8^, and are present in different stages of BC development and progression, as well as across BC molecular subtypes^9^. BCSCs may arise from differentiated breast cells or normal mammary stem cells and progenitors, owing to an acquired cell plasticity phenotype which closely associates with epithelial-to-mesenchymal transition (EMT) ^10-12^. In addition, BCSCs with different characteristics may be simultaneously present in the same tumor or in tumors of different BC molecular subtypes, including a mesenchymal-like BCSC subpopulation characterized by CD44+ CD24-EpCAM-marker expression ^13^. The high plasticity and phenotypic heterogeneity of BCSCs present challenges to their therapeutic targeting ^9^, highlighting the need to further explore BCSC states, their drivers and their specific stemness-associated phenotypes.

We previously observed that genetic and expression alterations in ERGs are frequent in BC patients ^3^, supporting the notion that ERG deregulation may contribute to BC development. These alterations may potentially underpin the epigenome alterations described in BC tumors, such as those leading to aberrant expression of CSC and EMT-associated genes ^14-16^. Still, most of the available data on epigenetic alterations in BC fail to capture contributions of specific ERGs to different steps of tumor development and progression, including the emergence of cells with BCSC properties. Moreover, specific epidrivers of cancer cell plasticity and BCSCs are yet to be identified and characterized in depth.

In this study, we used a powerful CRISPR/Cas9 screening tool combined with an *in-silico* approach focused on ERGs to identify potential epidrivers contributing to the emergence of a mesenchymal BCSC-like phenotype in non-tumorigenic mammary cells. We further employed mechanistic and multi-omics approaches for an in-depth characterization of a top candidate epidriver, the deubiquitinase BAP1. With this, we gained insight into the BAP1-mediated epigenetic regulation of different cellular processes and the acquisition of BC cell plasticity.

## METHODS

### CRISPR/Cas9 library screening

To identify ERG alterations contributing to a mesenchymal BCSC-like phenotype, we employed a CRISPR/Cas9 screening approach using a custom-made lentiviral library of 1,649 guide RNAs (gRNAs) targeting 426 ERGs ^3^, including writers, readers and editors of epigenetic marks such as DNA methylation, histone post-translational modifications and chromatin remodeling, as well as chromatin regulators. Screening was performed in duplicates (n=2) in two independent Cas9-expressing MCF10A clones (MCF10A-Cas9 clones 2 and 11). Details on gRNA library and generation of cell lines stably expressing Cas9 cells are provided in ^3^.

MCF10A-Cas9 clones were transduced with the lentiviral gRNA library at a multiplicity of infection (MOI) of 0.1 (day -6). Nontargeting gRNA LentiArray CRISPR Negative Control Lentivirus (Thermo Fisher Scientific) was used as a negative control. Two days after infection (day -4), cells were treated with puromycin (1 µg/mL) to select transduced cells until day 0, when non-transduced cells cultured in parallel were dead after selection. At least 2x10^6^ cells were maintained at each passage to ensure an average representation of 1,000x for each gRNA. Library-infected cells were sorted into different populations (BD FACSAria) according to cell surface marker expression on weeks 4 and 7 after puromycin selection for MCF10A clone 2 and clone 11 respectively, using antibodies for CD44 (1:400, 130-113-341, Miltenyi Biotech), CD24 (1:100, 130-112-656, Miltenyi Biotech) and EpCAM (1:200, 130-111-000, Miltenyi Biotech).

To identify enriched and depleted gRNAs in each cell population assessed, we isolated genomic DNA from 2x10^6^ cells (ensuring an average of 1 000x representation for each gRNA) with the AllPrep DNA/RNA Mini Kit (Qiagen), followed by PCR amplification of the gRNAs using forward (5’-CGATACAAGGCTGTTAGAGAGATA-3’) and reverse (5’-GTTGCTATTATGTCTACTATTCTTTCCC-3’) primers and NEBNext HighFidelity 2X PCR Master Mix (Illumina). Library preparation was performed with the Nextera DNA Flex Library Prep Kit (Illumina) according to manufacturer instructions and samples were sequenced on an Illumina MiSeq platform. FASTQ data generated by MiSeq were first analyzed in Galaxy using the BLASTN tool (v2.5.0) to obtain gRNA read counts, by mapping of identified gRNAs against all gRNA sequences in the CRISPR library. We next performed paired analysis comparing the gRNAs in the CD44^high^ EpCAM^-^ CD24^low^ sorted population with gRNAs present in the bulk of cells at the day of sorting, using clone 2 and clone 11 gRNA read counts as replicates allowing for more stringency. The differential analysis of the gRNA read counts was performed with CRISPRAnalyzeR (DKFZ, Version: 1.50) ^17^, using Model-based Analysis of Genome-wide CRISPR/Cas9 Knockout (MaGeCK) ^18^ (p-value threshold of < 0.05).

### Analysis of ERG mutations in breast cancer patient data

Breast cancer patient data from The Cancer Genome Atlas Breast Cancer (TCGA-BRCA) database was extracted from cBioPortal (https://www.cbioportal.org/datasets). According to the information on receptor expression provided by the database, a total of 1,097 samples were stratified based on the expression of ER, PR and HER2 into the following subtypes: 1) ER+HER2-, 588 cases; 2) ER+HER2+, 141 cases; 3) ER-HER2+, 41 cases; and 4) TNBC, with no expression of ER, PR or HER2, 158 cases. We excluded 169 samples with unknown or conflicting expression of ER and/or HER2. The proportion of single nucleotide alterations (SNAs) in ERGs was calculated by counting the number of individuals from each BC subtype (group) which carry at least one SNA for each gene and dividing this value by the total number of individuals in the group. We considered ERGs as frequently mutated when somatic mutations in a specific gene were identified in over 1% of BC patients, allowing the discovery of potential BC epidrivers which are less frequently altered than well-known BC driver genes such as *TP53*, *ERRB2* and *PI3KCA*. These genes, although mutated in over 10% of cases, represent only a small portion of potential BC drivers ^19-21^.

### Survival analysis

Transcriptomic data was obtained from TCGA-BRCA, using log-transformed RNA-seq values normalized by the FPKM method [log2(1+FPKM)]. To identify an optimal gene expression threshold for *BAP1* which stratifies individuals into “low” and “high” expression groups, we used the “ectopy” method as previously described ^22,23^, considering a range of possible thresholds from 10^th^ to 90^th^ percentile. Each threshold was tested for its ability to discriminate between two groups of tumors (of low and high expression values) corresponding to significantly different disease-free survival probabilities. All thresholds were analyzed using Cox proportional hazard model between two groups of low and high expression levels, and an optimal threshold with a significant p-value lower than 0.05 was selected. For the *BAP1* gene, the optimal threshold identified corresponds to the 10^th^ percentile of the total distribution of expression values in tumor samples. The software implemented for survival analysis is publicly available from https://github.com/epimed/ectopy.

### Generation of BAP1 knockout (KO) and rescue cells

MCF10A-Cas9 clone 11 cells were plated at a density of 5x10^4^ cells in 12-well plate wells, and transfected the next day with a pool of four BAP1 gRNAs at a final concentration of 2 µg (0.5 µg of each BAP1 gRNA), using Xfect Transfection Reagent (Takara Bio) according to the manufacturer’s instructions. gRNA sequences used are available in Supplementary Table S1. After 48h, transfected cells were selected with puromycin (1 µg/mL) until control non-transfected cells were dead. Individual KO clones were generated by sorting the bulk of antibiotic-selected cells into single cells in 96-well plates through flow cytometry (BD FACSAria), followed by amplification of the clones.

Wildtype and catalytically inactive (C91S) BAP1 plasmids were obtained from Addgene (#154020 and #154021) and packaged into retroviral particles using Phoenix cells and CalPhos Mammalian Transfection Kit (BD Biosciences), as previously described ^24^. MCF10A BAP1 KO cells were plated into 25 cm^2^ flasks at a density of 5x10^5^ cells, and infected the next day with 6mL of the collected and filtered Phoenix supernatant containing retroviral particles with either the wildtype or the mutant BAP1 construct. 24h after infection, cells were selected with 20 µg/mL of hygromycin until all non-infected control cells were dead.

Validation of KOs and rescues was carried out with western blot for BAP1 expression.

### ATAC-sequencing

Changes in chromatin accessibility were assessed with Assay for Transposase-Accessible Chromatin with high-throughput sequencing (ATAC-seq) in MCF10A-Cas9 and two clones of MCF10A-Cas9-BAP1 KO cells cultured as mammospheres, in duplicates (n=2), using the Omni-ATAC protocol as previously reported ^25^. Briefly, a 45-minute transposition step using the Tagment DNA TDE1 Enzyme (Illumina) was performed on 5x10^4^ viable cells per condition, followed by amplification with the NEBNext High-Fidelity 2X PCR Master Mix (New England Biolabs), using oligos available from ^26^. Library DNA cleanups were performed with the DNA Clean & Concentrator-5 Kit (Zymo Research). Libraries were sequenced as 150 bp paired-end reads on Illumina NextSeq 500, using the High Output Kit v2.5 (150 cycles, ref. 20024907).

### Chromatin immunoprecipitation (ChIP) and ChIP-sequencing

ChIP was performed in duplicates (n=2) in MCF10A wildtype (MCF10A-WT), MCF10A-Cas9 and two clones of MCF10A-Cas9-BAP1 KO cells cultured in attachment, using the iDeal ChIP-seq Kit for Histones (Diagenode) according to the manufacturer’s protocol. One million cells per IP were sonicated for 8 cycles [30 seconds “ON”, 30 seconds “OFF”] on a Bioruptor Pico (Diagenode), and immunoprecipitated with anti-H2AK119ub1 antibody (1:100 dilution; D27C4, Cell Signaling Technology). 10 ng of the H2AK119ub1 ChIPed DNA and inputs were used for library preparation with the TruSeq ChIP Sample Preparation Kit (Illumina), and samples were sequenced as 150 bp paired-end reads as described above.

### RNA-sequencing

RNA-sequencing (RNA-seq) was performed in duplicates (n=2) in both attachment- and mammosphere-grown MCF10A-Cas9 and two clones of MCF10A-Cas9-BAP1 KO cells, using an initial amount of 2 µg or 1 µg of total RNA for attached and mammosphere-cultured cells respectively. Libraries were prepared using the KAPA Stranded mRNA-Seq Kit (Kapa Biosystems) following the manufacturer’s protocol and sequenced as 75 bp single-end reads on Illumina NextSeq 500, with the High Output Kit v2.5 (75 cycles, ref. 20024906).

### Bioinformatic analyses of sequencing data

ATAC-seq reads were mapped to hg38 with BWA v0.7.17 ^27^ and trimmed using Trim Galore v0.6.10 ^28^. Peak signals were called with MACS2 ^29^, using the options: –extsize 150, –shift 75, and –keep-duplicates 1, and filtered using irreproducible discovery rate (IDR) within each condition. We considered the union of resulting peaks and called reads without considering duplicated ones. Differential analysis was performed with DiffBind v3.12.0 ^30^, by grouping the two BAP1 KO clones together and comparing changes with MCF10A-Cas9 controls (FDR < 0.01). Read numbers per peak were normalized using the RLE method, and differential peaks were called using DESeq2’s internal method, with a false discovery rate (FDR) cutoff of < 0.01. Differential peaks were annotated using a customized annotation pipeline that included gene coordinates from GENCODE V43 ^31^, information on driver genes using combined data from the COSMIC Cancer Gene Census ^32^ and the InTOGen database ^33^, 15 chromatin states for human mammary epithelial cells from the ChromHMM project ^34^ and Candidate cis-Regulatory Elements (CcREs) from ENCODE for mammary tumor cells (MCF-7) ^35^.

Estimation of differentially bound transcription factor motifs between BAP1 KOs and MCF10A-Cas9 control was performed with the TOBIAS package ^36^, using the BINDetect tool with standard parameters and motif annotations from the JASPAR2024 vertebrates database ^37^. BigWig signal files were inputted individually for each BAP1 KO clone and the MCF10A-Cas9 control, along with the list of ATAC-seq differential peaks (FDR < 0.01, annotated within 2kb upstream from TSSs). Transcription factors with differential binding score > |0.1| and p-value < 0.01 were considered significant. Results were plotted using the ggplot2 CRAN package, and Pearson correlation between replicates was calculated using the ggpurb CRAN package. In addition, a binned motif enrichment analysis approach was performed for the identification of differentially bound transcription factor motifs. This was achieved with the MOtif aNAlysis with Lisa (monaLisa) Bioconductor package ^38^ using the JASPAR2020 vertebrates database ^39^, with an FDR cutoff of 0.05.

ChIP-seq data was analyzed as described for ATAC-seq, with the –broad option in macs2 and considering duplicates for peak calling and read counting. Differential analysis was performed by comparing H2AK119ub1 peaks in BAP1 KOs with those identified in MCF10A-Cas9 and MCF10A-WT cells (FDR < 0.05). Custom annotation was performed as described above.

RNA-seq reads were mapped to hg38 with STAR v2.7.3a ^40^ and counted with HTSeq v0.12.4 ^41^. Read counts were normalized using median of ratios transformation with DESeq2 v1.38.3 ^42^. Both BAP1 KO clones were grouped together for differential expression analysis, which was also performed with DESeq2. Genes were considered as differentially expressed between comparisons if FDR < 0.01.

### Gene set and gene ontology enrichment analyses

Gene set enrichment analysis (GSEA) was performed on normalized gene counts from RNA-seq with the GSEA v4.3.2 software ^43^, using the hallmark gene set collection from the Molecular Signatures Database (MSigDB) ^44^ and an FDR cut-off of 0.05, after filtering out low count measurements from RNA-seq (<100 counts). Gene ontology (GO) enrichment analysis of biological processes was performed with clusterProfiler v4.2.2 ^45^ in R (v4.1.2) on differentially expressed genes from RNA-seq (FDR < 0.01) and the genes annotated closest to differential ATAC-seq peaks (FDR < 0.01), as well as on genes altered across different omics analyses. The “simplify” function was used to generate specific GO enrichment plots (Figures 3D and 4B, Supplementary Figure S4C), to replace redundant GO terms with a representative term (similarity cutoff of 0.5, i.e., GO terms with over 50% similarity are replaced by a representative term).

### Total proteomics analysis

Total proteomics analysis was performed in triplicates (n=3) in mammosphere-grown MCF10A-Cas9 and MCF10A-Cas9-BAP1 KO Cl1, using a total of 2 million cells per replicate.

*Sample preparation.* Cell pellets were collected and washed 2x with PBS, then lysed in Lysis Buffer (2% SDS, 50mM Tris pH 8.5, 10mM TCEP, 40mM CAA, supplemented with protease inhibitor cocktail and phosphatase inhibitors), sonicated at 4°C in a Bioruptor Pico2 (30/30, 10 cycles) and boiled at 95°C for 10 min. Proteins were precipitated using methanol-chloroform and resuspended in 8M urea, 50mM Tris pH 8.5. Proteins were digested with 1:50 w/w LysC (Wako Chemicals) and 1:100 w/w trypsin (Promega) overnight at 37°C after dilution to a final urea concentration of 1M using 50mM Tris pH 8.5. Digested peptides were then acidified (pH 2–3) using trifluoroacetic acid (TFA) and purified using C18 SepPak columns (Waters). Desalted peptides were dried and resuspended in TMT-labeling buffer (200mM EPPS pH 8.2, 20% acetonitrile). 10μg of peptides per condition were subjected to TMT labeling with 1:2.5 peptide TMT ratio (w/w) for 1 hour at room temperature. The labeling reaction was quenched by addition of hydroxylamine to a final concentration of 0.5% and incubation at room temperature for 15 min. Successful TMT labeling was verified by mixing equimolar ratios of peptides and subjecting the mix to single shot LC-MS/MS analysis. Peptides were fractionated using high-pH liquid-chromatography on a micro-flow HPLC (Dionex U3000 RSLC, Thermo Scientific). 45µg of pooled and purified TMT labelled peptides resuspended in Solvent A (5mM ammonium-bicarbonate, 5% ACN) were separated on a C18 column (XSelect CSH, 1mm x 150mm, 3.5µm particle size; Waters) using a multistep gradient from 3-60% Solvent B (5mM ammonium-bicarbonate, 90% ACN) over 65 minutes at a flow rate of 30µl/min. Eluting peptides were collected every 43 seconds from minute 2 for 69 minutes into a total of 96 fractions, which were cross-concatenated into 24 fractions. Pooled fractions were dried in a vacuum concentrator and resuspended in 2% ACN, 0.1% TFA for LC-MS analysis.

*Mass spectrometry data acquisition.* Tryptic peptides were analyzed on an Orbitrap Ascend coupled to a VanquishNeo (ThermoFisher Scientific) using a 25cm long, 75µm ID fused-silica column packed in house with 1.9µm C18 particles (Reprosil pur, Dr. Maisch), and kept at 50°C using an integrated column oven (Sonation). HPLC solvents consisted of 0.1% Formic acid in water (Buffer A) and 0.1% Formic acid, 80% acetonitrile in water (Buffer B). Assuming equal amounts in each fraction, 400ng of peptides were eluted by a non-linear gradient from 8% to 32% Buffer B over 77 minutes followed by a stepwise increase to 95% Buffer B in 11 minutes which was held for another 7 minutes. A synchronous precursor selection (SPS) multi-notch MS3 method was used in order to minimize ratio compression as previously described ^46^. Full scan MS spectra (350-1,400m/z) were acquired with a resolution of 120,000 at m/z 200, maximum injection time of 100ms and AGC target value of 4x10^5^. The most intense precursors with a charge state between 2 and 5 per full scan were selected for fragmentation (“Top Speed” with a cycle time of 1.5 seconds) and isolated with a quadrupole isolation window of 0.7 Th. MS2 scans were performed in the Ion trap (Turbo) using a maximum injection time of 35ms, AGC target value of 10,000 and fragmented using CID with a normalized collision energy (NCE) of 35%. SPS-MS3 scans for quantification were triggered only after successful Real-time search against the human canonical reference proteome from SwissProt with the same search parameter as stated below for data processing in Proteome Discoverer. Criteria for passing the search were Xcorr: 1, dCn: 0.1 and precursor mass accuracy: 10ppm. Maximum search time was 35ms. MS3 acquisition was performed on the 10 most intense MS2 fragment ions with an isolation window of 0.7Th (MS) and 2m/z (MS2). Ions were fragmented using HCD with an NCE of 50% and analyzed in the Orbitrap with a resolution of 45,000 at m/z 200, scan range of 100-200m/z, AGC target value of 150,000 and a maximum injection time of 91ms. Repeated sequencing of already acquired precursors was limited by setting a dynamic exclusion of 60 seconds and 7ppm and advanced peak determination was deactivated. All spectra were acquired in centroid mode.

*Data analysis.* MS raw data were analyzed using FragPipe v21.1, with MSFragger v.4.0 ^47^ and Philosopher v.5.1.0 ^48^. Acquired spectra were searched against the human reference proteome (Taxonomy ID 9606) downloaded from UniProt (07/25/2024; 20,418 entries) with a precursor mass tolerance of 20ppm and fragment mass tolerance of 20ppm. Identifications were filtered to obtain false discovery rates (FDR) below 1% for both peptide spectrum matches (minimum peptide length of 7) and proteins using a target-decoy strategy. For all searches, carbamidomethylated cysteine was set as a fixed modification and oxidation of methionine and N-terminal protein acetylation as variable modifications with allowing up to 3 modifications per peptide. Strict trypsin cleavage was set as protein digestion rule. Label-free quantification was performed using IonQuant v.1.10.27 ^49^. Data were further processed using FragPipe Analyst ^50^.

Briefly, following cell lysis, total proteins were precipitated and digested, and resulting peptides were labelled with TMT buffer (200 mM EPPS pH 8.2, 20% acetonitrile). Mass spectrometry data was acquired on an Orbitrap Ascend coupled to a VanquishNeo (ThermoFisher Scientific), and identified peptides were mapped to the human reference proteome (ID 9606). Proteins were considered as differentially expressed if FDR < 0.01.

### Glycan profiling

*N*-glycan profiling of MCF10A-Cas9 and MCF10A BAP1 KO cells cultured as mammospheres was performed with 1 million cells per replicate, for a total of 4 replicates per condition (n=4).

*Cell lysis, protein denaturation, alkylation, and enzymatic release of N-glycans*. Cell pellets of MCF10A-Cas9 and MCF10A BAP1 KO cells cultured as mammospheres, containing 1 million cells each, were resuspended in 0.2% solution of RapiGest SF surfactant (Waters) in 1x PBS, with 5 µL 60 mM DTT (Sigma-Aldrich). Samples were sonicated for 5 minutes in an ultrasonic bath, followed by protein denaturation for 15 minutes at 60°C. After cooling down, 4 µL 160 mM IAA (Sigma-Aldrich) solution was added and protein alkylation was carried for 30 minutes, protected from light. Excess IAA was quenched by adding 1 µL of DTT solution. For deglycosylation, 1 U of PNGase F (Promega) in 5 µL 1x PBS was added and samples were incubated overnight at 37°C.

*Fluorescent N-glycan labelling and purification*. Released *N*-glycans were labelled with procainamide (ProA, Acros Organics) via reductive amination. First, 4.32 mg ProA dissolved in 25 µL 30% glacial acetic acid in DMSO (*v*/*v*) was added to each sample and incubated at 65°C. Then, 4.48 mg of 2-picoline borane complex (Sigma-Aldrich) in 25 µL of the same solvent was added and incubated for another 1.5 hour. After cooling down, 1,080 µL of cold ACN was added and ProA-labelled N-glycans were purified first via Hydrophilic Interaction Liquid Chromatography Solid-Phase Extraction (HILIC-SPE) ^51^ and then on Porous Graphitic Carbon (PGC) as previously described ^52^.

*N-glycan analysis*. For HILIC-Ultra-performance liquid chromatography with fluorescence detection (HILIC-UPLC-FLR) analysis of N-glycans, we employed the Waters H-class UPLC system equipped with quaternary pump, sample manager and fluorescence detector (λ_ex_/λ_em_ = 310/370 nm). ProA-labelled N-glycans (40 µL) were injected and separated on a Waters UPLC glycan bridged ethylene hybrid (BEH) amide chromatographic column (150 × 2.1 mm, 1.7 µm), with 100 mM ammonium formate (pH 4.4) as mobile phase A, and acetonitrile as mobile phase B. The separation method used a linear gradient of 25 to 40 % mobile phase A at the flow rate of 0.56 mL/minute in a 98-minute analytical run. Samples were maintained at 10°C before injection, and separated at 50°C. UPLC system was operated under Empower software (v3.6.1). The system was calibrated using an external standard of hydrolyzed and ProA-labelled glucose oligomers, from which the retention times for the individual N-glycans were converted to glucose units. Data processing was performed using an automatic processing method with a traditional integration algorithm, after which each chromatogram was manually corrected to maintain the same intervals of integration for all the samples. The chromatograms were all separated in the same manner into 47 peaks, and the amount of N-glycans in each peak was expressed as % of total integrated area.

### Visualization and statistical analyses

GraphPad Prism v9.5.1 and v10.0.0 were used for the preparation of plots and statistical analyses of RT-qPCR, cell surface protein expression, pyrosequencing (plots only), proliferation, cell cycle, and metabolic assays. R v4.1.2 was used for volcano plots. ATAC-seq and H2AK119ub1 peaks were visualized on IGV (version 2.12.3), using bigwig and bed files.

Description of additional methods is available from the Supplementary Information file.

## RESULTS

### CRISPR/Cas9 loss-of-function screening identifies putative epidrivers of mesenchymal breast cancer stem cells

To identify key epidrivers in the emergence of mammary cells with mesenchymal-like BCSC characteristics, we employed a lentiviral CRISPR/Cas9 loss-of-function (LOF) screening approach targeting 426 ERGs in non-tumorigenic MCF10A mammary cells stably expressing Cas9 (MCF10A-Cas9) (Figure 1A). A mesenchymal BCSC-like population (CD44^high^ CD24^low^ EpCAM^-^) was isolated from the library-infected cells, and enriched gRNAs were identified through MiSeq sequencing. We identified six ERGs whose LOF could be linked to the emergence of a mesenchymal BCSC-like phenotype (p-value < 0.05), all associated with histone post-translational modification: *ASXL2*, *BAP1*, *CBX4*, *KAT6B*, *KDM6A* and *PRDM9* (Figure 1B, Supplementary Table S4). Among these, the *BAP1* and *ASXL2* genes, forming the Polycomb repressive deubiquitinase complex (PR-DUB), and the *KDM6A* gene, part of the COMPASS complex (complex of proteins associated with SET1), were previously described to regulate a permissive chromatin landscape ^53,54^.

**Figure 1.**
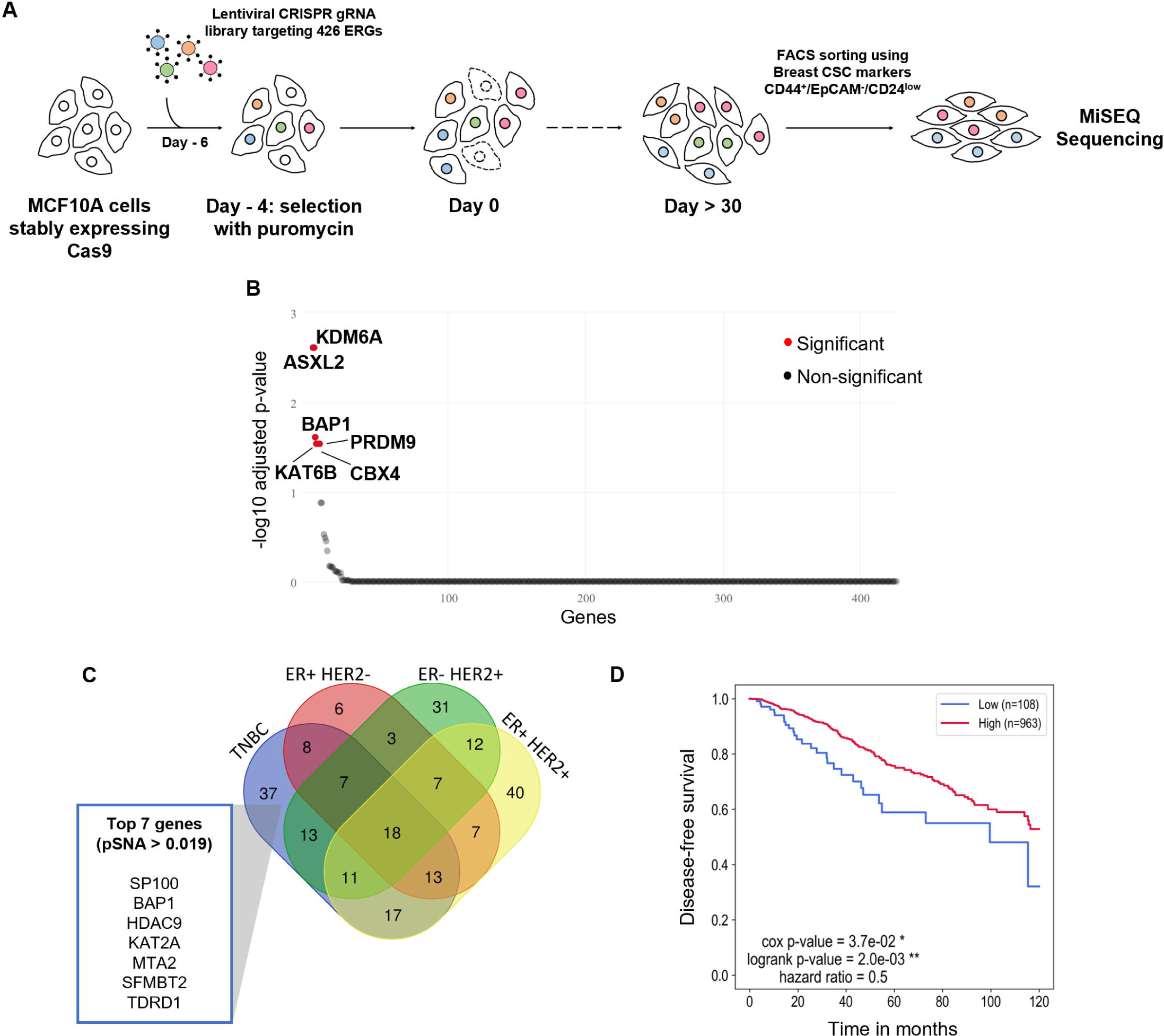
BAP1 disruption is a putative epidriver of mesenchymal breast cancer stem cell-like cells. (A) CRISPR/Cas9 screening approach used for the identification of epigenetic regulator genes (ERGs) involved in the acquisition of mesenchymal breast cancer stem cell (BCSC) markers in non-tumorigenic breast cells. Adapted from Halaburkova et al. 2020 ^3^. gRNA: guide RNA. (B) Representation of enriched ERG gRNAs (false discovery rate [FDR] < 0.05) identified in the mesenchymal BCSC-like population of MCF10A cells infected with the ERG gRNA library compared to the bulk of cells on the day of sorting (n=2 MCF10A-Cas9 expressing clones). (C) Venn diagram of ERGs showing single nucleotide alterations in BC patients (TCGA-BRCA) of different molecular subtypes. Top 7 mutated ERGs identified in the TNBC subtype (proportion of SNAs (pSNA) > 0.019) are highlighted. (D) Kaplan-Meier analysis of disease-free survival in BC patients (TCGA-BRCA) divided in high and low BAP1 gene expression groups. * *P* < 0.05, ** *P* < 0.01.

Interestingly, *ASXL2*, *BAP1* and *PRDM9* were among the topmost frequently mutated ERGs in BC patients (Figure 1C, Supplementary Table S5). Of note, *BAP1* and *PRDM9* were exclusively altered in triple negative breast cancers (TNBCs), suggesting a potential role in this cancer subtype, characterized by an enrichment in highly plastic cells and elevated expression of BCSC markers ^55,56^. Assessment of the clinical impact of ERG expression in BC patients revealed that diminished expression of *BAP1* correlates with poorer disease-free survival (Figure 1D), pointing to *BAP1* disruption as a possible prognostic marker in BCs. Based on these findings, we hypothesized that loss of *BAP1* function could be an epidriver of the emergence of mesenchymal BCSC-like cells in BC development. *BAP1* is described as a tumor suppressor, and both germline and somatic mutations in this gene have been reported in different cancer types ^57,58^. However, little is known regarding the mechanistic contributions of *BAP1* deregulation to cancer cell plasticity and tumor biology in BC.

### BAP1 loss in non-tumorigenic breast cells enhances expression of BCSC and EMT markers

To validate the role of *BAP1* as a putative epidriver of breast cancer cell plasticity and the emergence of mesenchymal BCSC-like markers in mammary cells, we generated two independent MCF10A-Cas9 clones lacking BAP1 expression (BAP1 knockouts, KOs) (Figure 2A). When grown as mammospheres (mammary spheroids), a three-dimensional (3D) culture condition which allows enrichment of breast (cancer) stem cells ^59^, both BAP1 KO clones exhibited visible changes in spheroid morphology (Figure 2B). While control MCF10A-Cas9 mammospheres displayed a round and compact shape, BAP1 KO spheroids exhibited a diffused and less uniform surface suggestive of a loss of contact between neighboring cells, a characteristic of mesenchymal cells. BAP1 KO mammospheres showed slightly reduced cell adhesion in their periphery, represented by detached cells at the borders, and presence of some peripheral bodies and intracytoplasmic vacuoles. These changes may be associated with disruptions in cell-to-cell communication and/or cell migration ^60^ (Figure 2C, Supplementary Figure S1), in line with previous reports of alterations in the organization of three-dimensional cellular structures upon BAP1 loss ^61,62^.

**Figure 2.**
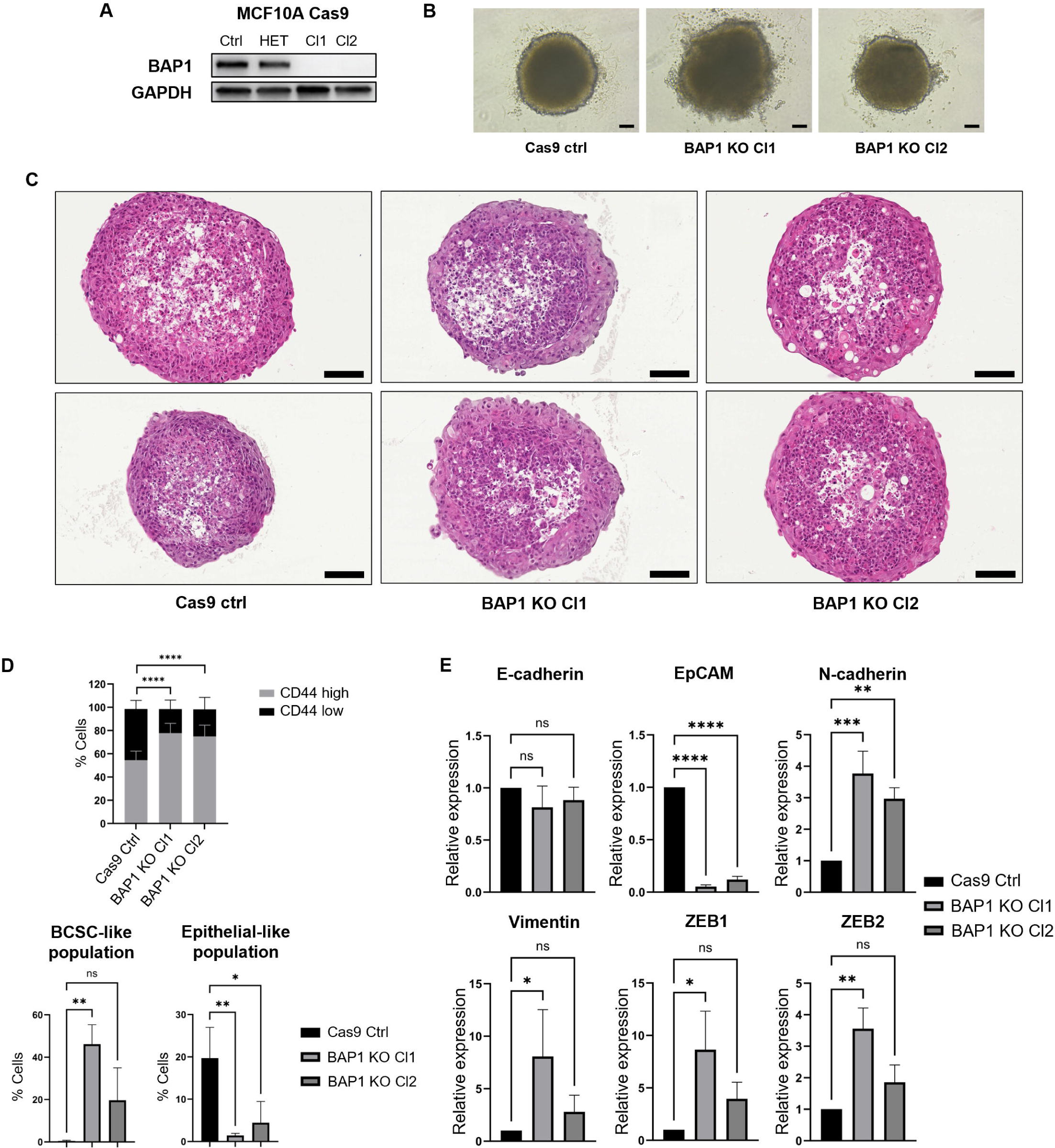
BAP1 loss promotes changes in mammosphere architecture and enhances expression of BCSC and EMT markers. (A) BAP1 protein expression levels in MCF10A-Cas9 control cells (Ctrl), a heterozygous BAP1 deletion clone (HET), and two BAP1 knockout (KO) clones (Cl1 and Cl2). GAPDH levels were assessed as an internal control. (B) Representative images at 10x magnification of single, size-normalized MCF10A-Cas9 control and BAP1 KO mammospheres cultured for 72h. Scale bars: 100 µm. (C) Representative images at 20x magnification of single, size-normalized MCF10A-Cas9 control and BAP1 KO mammospheres stained with hematoxylin and eosin show mammosphere architecture differences and cellular changes between BAP1 KOs and controls. BAP1 KOs show disorganized architecture and a variable amount of intracytoplasmic vacuolization and cellular decohesion, all noted at the periphery of the mammospheres. Scale bars: 100 µm. See also Supplementary Figure S1. (D) Top: distribution of CD44 high and low populations in MCF10A-Cas9 controls and BAP1 KO cells cultured as mammospheres. Significance analysis performed by Fisher’s exact test on absolute cell counts (n=3, **** *P* < 0.0001). Bottom: proportion of BCSC- and epithelial-like populations based on expression of CD44, CD24 and EpCAM in MCF10A-Cas9 controls and BAP1 KO cells cultured as mammospheres. Significance analysis performed by Dunnett’s test (n=3, * *P* < 0.05, ** *P* < 0.01, ns: not significant). (E) Expression of EMT-associated genes in MCF10A-Cas9 and BAP1 KO cells cultured as mammospheres. Significance analysis performed by Dunnett’s test (n=3, * *P* < 0.05, ** *P* < 0.01, *** *P* < 0.001, **** *P* < 0.0001, ns: not significant).

Next, we confirmed that BAP1 loss leads to the acquisition of mesenchymal BCSC-like markers in mammospheres, with a significant increase in CD44 expression and a striking shift in the proportion of CD44^high^ EpCAM^-^ CD24^low^ mesenchymal BCSC-like cells (0.47% of cells in controls, 46.2% and 19.69% in BAP1 KOs), accompanied by a decrease in the CD44^low^ EpCAM^+^ CD24^high^ epithelial-like population (19.7% of cells in controls, 1.43% and 4.43% in BAP1 KOs) (Figure 2D, Supplementary Figure S2A). In addition, BAP1 KOs displayed changes in expression of EMT-associated genes, including decreased transcription of epithelial markers E-cadherin and EpCAM, and increased expression of mesenchymal markers N-cadherin, Vimentin, ZEB1 and ZEB2 (Figure 2E, Supplementary Figure S2B), suggesting a role of BAP1 loss in regulating the EMT program. In line with changes in gene expression, CpG islands in the promoter regions of the E-cadherin and ZEB1 genes showed consistent differences in DNA methylation between BAP1 KOs and MCF10A-Cas9 controls, especially in mammosphere-cultured cells (Supplementary Figure S2C and D). Together, these results indicate that BAP1 loss can promote the acquisition of BCSC-like and mesenchymal features in non-tumorigenic mammary cells.

### BAP1 loss leads to global reduction in chromatin accessibility associated with H2AK119ub1 accumulation

BAP1 is a deubiquitinating enzyme and the catalytical subunit of the PR-DUB complex, which regulates gene expression by removing the silencing histone mark H2AK119ub1 ^63^. Because removal of H2AK119ub1 by BAP1 prevents aberrant chromatin compaction, antagonizes Polycomb-mediated gene silencing and promotes transcription ^63^, we sought to explore the impact of BAP1 loss in the chromatin accessibility of non-tumorigenic breast cells by performing ATAC-seq in mammosphere-cultured BAP1 KOs and MCF10A-Cas9 controls. We observed a clear shift in the cells’ epigenetic state following BAP1 loss (Supplementary Figure S3A), with chromatin accessibility changes observed in 36,478 sites (FDR < 0.01), which we defined as either ATAC loss (reduced accessibility, log2FC < 0) or ATAC gain (increased accessibility, log2FC > 0) (Figure 3A). Over 77% of differential peaks showed ATAC loss in the KOs (n=28,345), suggesting widespread reduction in chromatin accessibility following BAP1 disruption. Interestingly, a prominent fraction of the less accessible peaks upon BAP1 disruption was found in promoter regions (15.7% of ATAC loss, 3.1% of ATAC gain) (Figure 3B, Supplementary Table S6), active transcription start sites (TSSs) (13.1% of ATAC loss, 0.5% of ATAC gain) and regions flanking TSSs (6.1% of ATAC loss, 3.5% of ATAC gain) (Figure 3C, Supplementary Table S7), suggesting that loss of accessibility in these peaks might have a direct impact on the transcription of associated genes.

**Figure 3.**
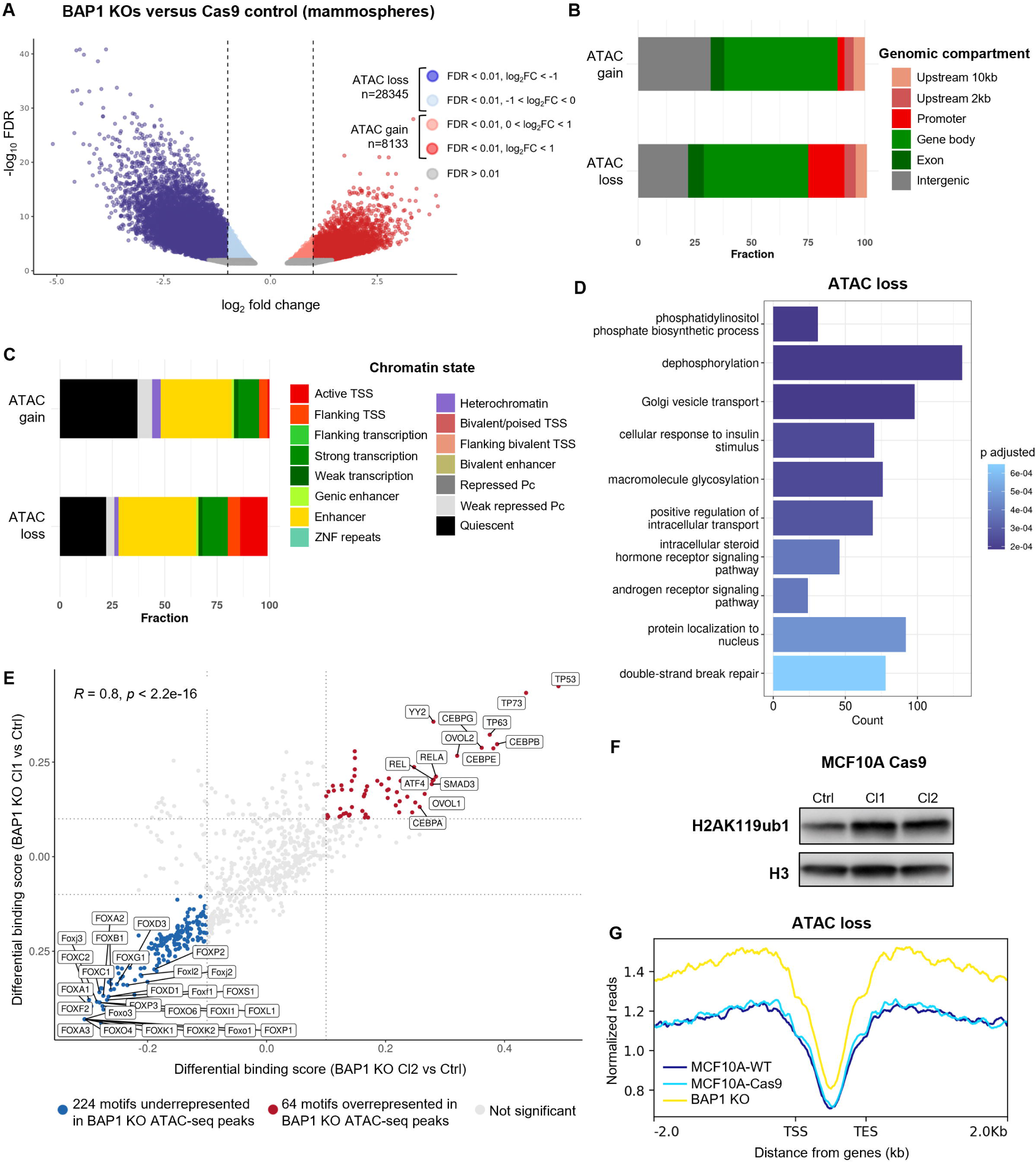
BAP1 loss leads to global reduction in chromatin accessibility and H2AK119ub accumulation in mammary cells. (A) Volcano plot of ATAC-seq peaks identified in BAP1 KOs (n=4) compared to MCF10A-Cas9 controls (n=2) cultured as mammospheres. Blue and red-colored dots represent peaks with loss and gain of ATAC-seq accessibility (log2FC < 0 and log2FC > 0), respectively. (B) Genomic compartment and (C) chromatin state distribution of differential ATAC-seq peaks (FDR < 0.01) identified in BAP1 KOs compared to MCF10A-Cas9 cells cultured as mammospheres. Promoters are defined as regions 500 base pairs (bp) upstream of transcription start sites (TSSs). Kb: kilobases. Pc: Polycomb. (D) Gene ontology enrichment of genes associated with differential ATAC loss peaks (FDR < 0.01, log2FC < 0) falling 2kb upstream from TSSs in BAP1 KOs compared to MCF10A-Cas9 cells cultured as mammospheres. Top 10 enriched ontologies are shown (p adjusted < 0.05). (E) Differential transcription factor binding in BAP1 KO clones compared to MCF10A-Cas9 (Ctrl) cells cultured as mammospheres. Blue: 224 underrepresented motifs (differential binding score < -0.1, p < 0.01); red: 64 overrepresented motifs (differential binding score > 0.1, p < 0.01); and gray: motifs with no representative alteration (-0.1 ≤ differential binding score ≤ 0.1) in ATAC-seq peaks from BAP1 KOs. (F) H2AK119ub1 levels in MCF10A-Cas9 control cells (Ctrl) and two BAP1 knockout (KO) clones (Cl1 and Cl2) cultured in attachment. H3 levels were assessed as an internal control. (G) Distribution of H2AK119ub1 ChIP-seq peaks (attached cells, FDR < 0.05) in regions of ATAC loss (mammosphere cultures, FDR < 0.01, log2FC < 0) identified in BAP1 KOs compared to MCF10A-Cas9 cells. Regions within 2kb upstream and downstream of TSSs and transcription end sites (TESs), respectively, are shown. ChIP-seq reads were normalized with RPGC (reads per genome coverage) per bin method. ChIP-seq peak distribution was analyzed for MCF10A wildtype (MCF10A-WT, n=2), MCF10A-Cas9 controls (MCF10A-Cas9, n=2), and both BAP1 KO clones (BAP1 KO, n=4).

Gene ontology (GO) analysis of genes associated with differential ATAC-seq peaks found within 2kb upstream of TSSs revealed reduced accessibility in genes involved in glycosylation, phospholipid synthesis and receptor signaling-associated processes (Figure 3D), suggesting that BAP1 may regulate cell metabolism and cellular membrane pathways. In contrast, accessible ATAC-seq peaks showed enrichment of fewer biological processes (Supplementary Figure S3B), which might be explained by the lower number of open peaks identified in regions 2kb upstream of TSSs following BAP1 loss.

Next, we investigated the enrichment of transcription factor binding motifs in differentially accessible chromatin regions within 2kb upstream of TSSs. We observed a high correlation in motif enrichment between both KO clones (R=0.8, p < 2.2 x 10^-16^) and a total of 288 motifs with significant enrichment (differential binding score > |0.1|, p < 0.01) in MCF10A BAP1 KOs compared to controls (Figure 3E, Supplementary Table S8). Most of the identified motifs were underrepresented in BAP1 KOs (224 motifs, differential binding score < -0.1), suggesting that BAP1 loss affects the binding of different transcription factors. Motifs with the lowest binding score in BAP1 KOs mainly include FOX transcription factors such as FOXK2, which was previously described to recruit BAP1 to chromatin ^64^. On the other hand, top overrepresented motifs in BAP1 KOs (64 motifs, differential binding score > 0.1) include the TP53 and TP73 proteins, a finding confirmed with a second motif analysis tool (Supplementary Figure S3C). Together, these results point to extensive chromatin remodeling and an overall reduction in chromatin accessibility following BAP1 loss, which may impact different cellular pathways and the binding of transcription factors.

We further assessed H2AK119ub1 levels in MCF10A cells following BAP1 disruption, and found an accumulation of this silencing mark upon BAP1 loss (Figure 3F). ChIP-seq for H2AK119ub1 revealed an overall gain in this histone mark in regions surrounding coding genes in BAP1 KOs, especially in regions overlapping with loss in chromatin accessibility (Figure 3G, Supplementary Figure S3D). Next, we annotated enriched H2AK119ub1 peaks (FDR < 0.05) to their closest associated gene, and found that approximately 16.4% of H2AK119ub1-enriched genes were associated with ATAC loss within 2kb upstream of their TSSs, whereas only 2.6% of these genes overlapped with ATAC gain (Supplementary Figure S3E) in BAP1 KOs. Together, these data reveal that BAP1 loss results in increased levels of H2AK119ub1, which associate with loss of accessibility in regions adjacent to gene transcription, suggesting that BAP1 may directly regulate chromatin accessibility in these regions through its histone deubiquitinase function.

### BAP1 loss deregulates expression of genes related to metabolic and glycosylation pathways

To explore the impact of BAP1 loss in gene transcription, we employed an RNA-seq approach which revealed distinct gene expression profiles between mammosphere-cultured BAP1 KOs and MCF10A-Cas9 controls (Supplementary Figure S4A). Upon BAP1 depletion, 3,925 differentially expressed genes (DEGs) (FDR < 0.01) were identified, most of them downregulated (log2FC < 0; 2,357 genes, approximately 60% of DEGs) (Figure 4A), in line with the overall loss in chromatin accessibility and the accumulation of H2AK119ub1 observed in BAP1 KOs. In attached cells, BAP1 KOs also exhibited a consistent downregulation in gene expression (1,536 genes out of 2,405, or 64% of DEGs) (Supplementary Figure S4B and S4C), supporting the notion that BAP1 loss leads to an overall downregulation of transcription in non-tumorigenic mammary cells.

**Figure 4.**
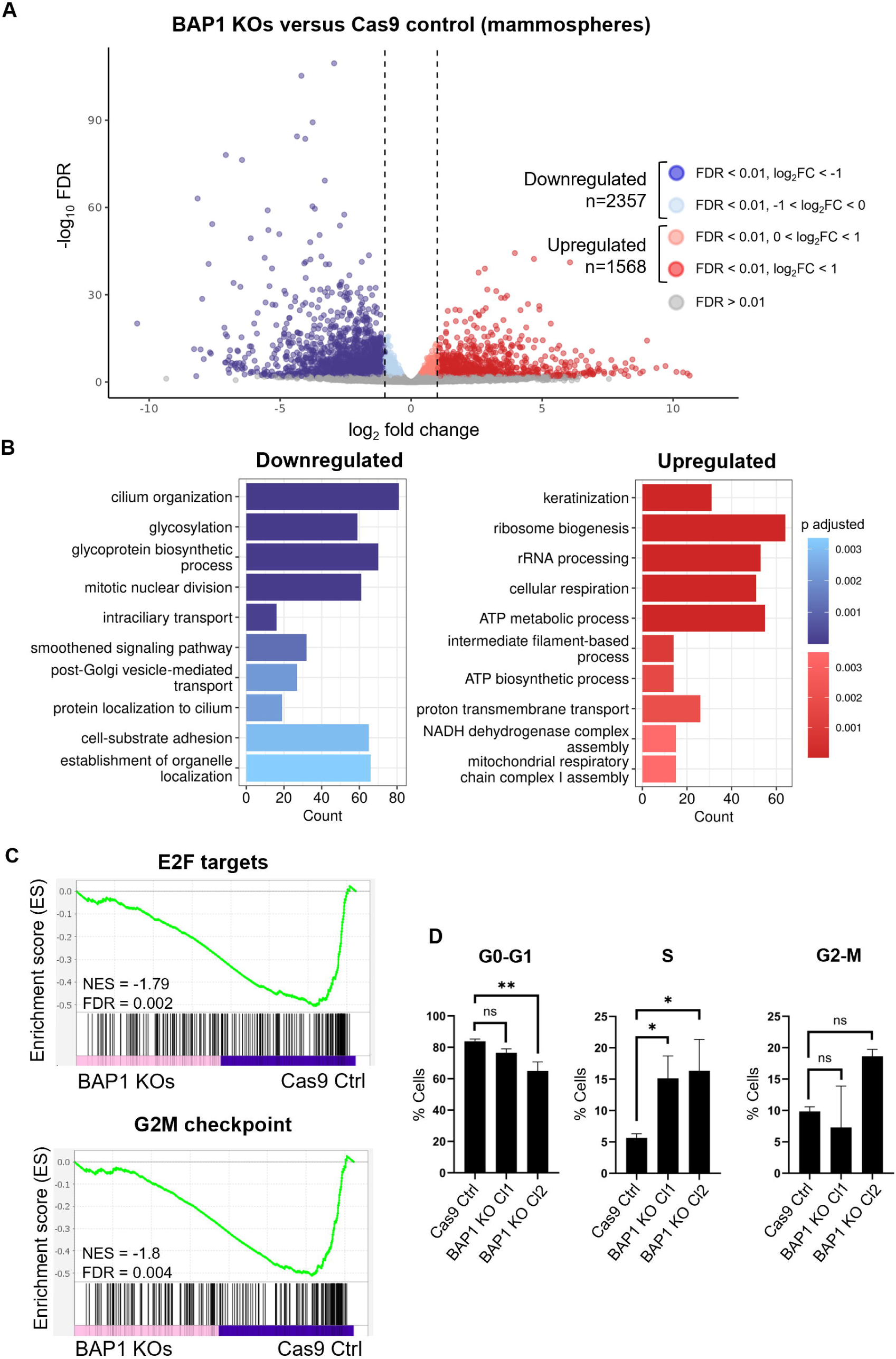
BAP1 loss deregulates gene expression of cell cycle, metabolic and glycosylation genes. (A) Volcano plot of differential expression analysis in BAP1 KOs compared to MCF10A-Cas9 control cells cultured as mammospheres (n=2 for MCF10A-Cas9 control cells and n=4 for BAP1 KOs). Blue and red-colored dots represent differentially expressed genes (DEGs, FDR < 0.01) with reduced or increased expression (log2FC < 0 and log2FC > 0), respectively. (B) Gene ontology enrichment of downregulated (left) and upregulated (right) DEGs (FDR < 0.01) in BAP1 KOs compared to MCF10A-Cas9 cells cultured as mammospheres. Top 10 enriched gene ontologies are shown (p adjusted < 0.05). (C) Gene set enrichment analysis (GSEA) plots of the transcriptome of BAP1 KOs compared to MCF10A-Cas9 control cells cultured as mammospheres, using the indicated gene sets. NES: normalized enrichment score. (D) Cell cycle analysis by propidium iodide staining in unsynchronized MCF10A-Cas9 and BAP1 KO mammospheres. Each bar graph shows the percentage distribution of cells in each cell cycle phase: G0-G1, S and G2-M. Significance analysis performed by Dunnett’s test (n=3, * *P* < 0.05, ** *P* < 0.01, ns: not significant).

GO analysis of DEGs revealed the upregulation of ATP metabolism and mitochondrial respiration pathways in mammosphere-cultured BAP1 KOs (Figure 4B), indicating a possible role in the control of cellular respiration and metabolic activity. However, an assessment of mammosphere metabolism showed no alterations in mitochondrial respiration and only a slight reduction in glycolytic capacity in BAP1 KOs (Supplementary Figure S5A and B).

Top downregulated biological processes upon BAP1 loss included cilium-associated and glycosylation pathways, which were also identified in cells cultured in attachment (Figure 4B, Supplementary Figure S4C and S6). Notably, glycosylation was among the top biological processes associated with ATAC loss in BAP1 KO mammospheres, indicating that changes in expression of glycosylation genes might result from loss in chromatin accessibility. In addition, we observed downregulation of cell cycle and mitosis-associated pathways in BAP1 KO mammospheres (Figure 4B). Similarly, gene set enrichment analysis (GSEA) revealed negative enrichment of the E2F targets and G2M checkpoint gene sets in BAP1 KOs, both involved in DNA replication and cell cycle progression ^44^ (Figure 4C), suggesting that BAP1 loss may impact normal cell cycle progression in mammospheres, potentially coupled with EMT. Indeed, we observed an accumulation of BAP1 KO cells in the S phase accompanied by lower levels of cells in G0-G1 (Figure 4D), and a tendency of reduced proliferation in MCF10A BAP1 KOs cultured in attachment compared to controls (Supplementary Figure S5C), in accordance with a previous report in this same cell line ^65^. Interestingly, different genes involved in cell division regulation showed reduced ATAC accessibility within 2kb upstream of their TSSs and/or associated with significant H2AK119ub1 gain in their vicinity in BAP1 KOs (Supplementary Figure S5D), suggesting that BAP1 may be involved in the transcriptional control of cell division genes through its epigenetic roles. Collectively, our transcriptomic analysis points to BAP1 as a regulator of gene expression programs associated with cell cycle progression, proliferation, glycosylation, and metabolism in non-tumorigenic mammary cells.

### Deregulation in gene expression following BAP1 loss is linked to BAP1’s functions as an epigenetic regulator

To assess whether the changes in gene expression following BAP1 loss are associated with its role as an epigenetic regulator, we overlapped DEGs identified in BAP1 KOs with differentially accessible ATAC-seq peaks within 2kb upstream of TSSs. This revealed that reduced chromatin accessibility generally associated with downregulated gene expression, with nearly half of the downregulated genes showing concomitant loss in chromatin accessibility in BAP1 KOs, whereas gene upregulation showed less overlap with ATAC-seq signal (Figure 5A, Supplementary Figure S7A). Importantly, the absence of changes in chromatin accessibility at most of the upregulated genes and approximately half of the downregulated genes (neutral ATAC-seq signal) may point to indirect or epigenetic regulation-independent effects of BAP1 loss.

**Figure 5.**
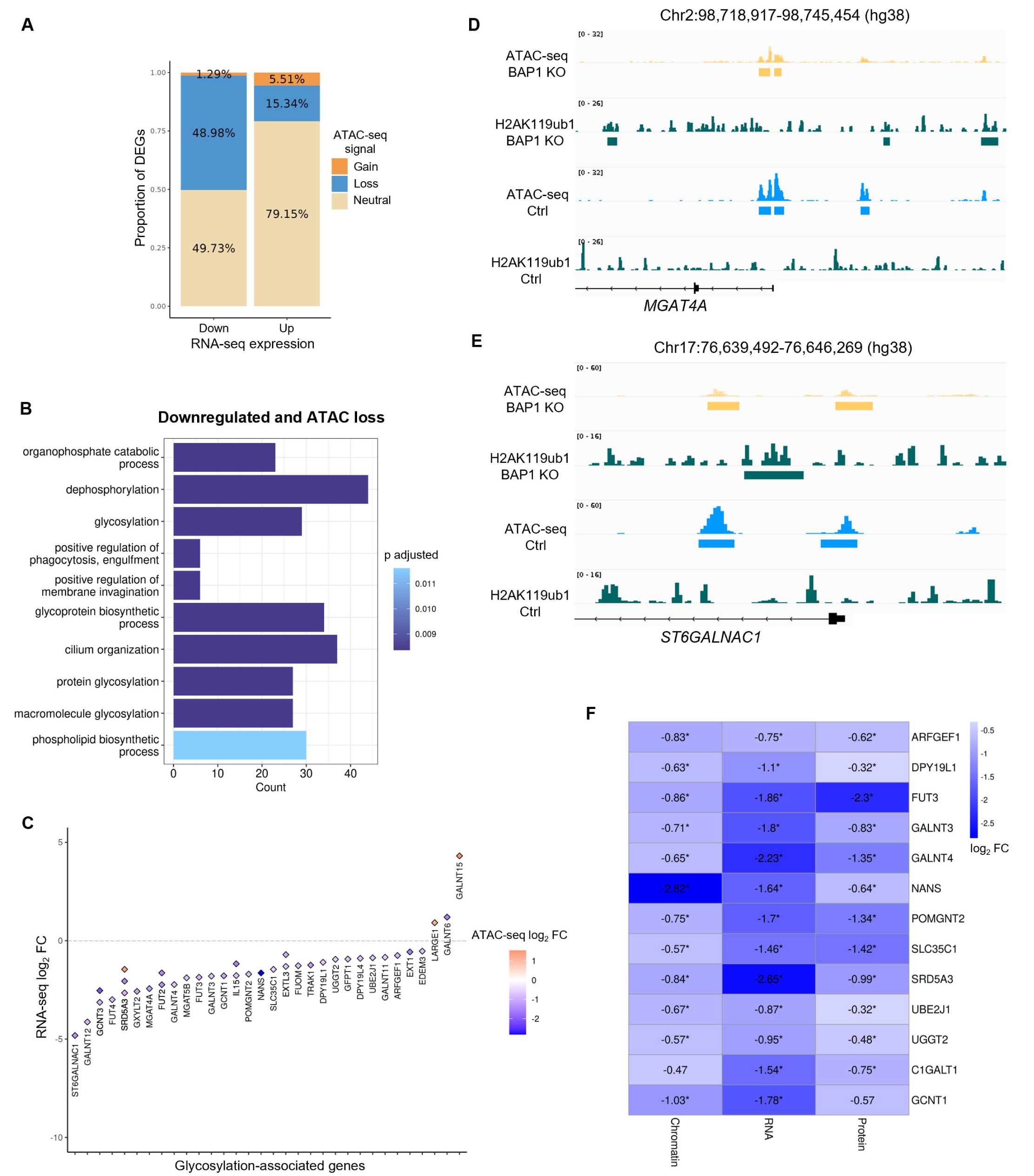
BAP1 loss-associated epigenetic deregulation impacts gene and protein expression. (A) Proportion of DEGs (FDR < 0.01) showing differential (FDR < 0.01) ATAC-seq loss (log_2_FC < 0), gain (log_2_FC > 0) or neutral signal (no loss or gain) within 2kb upstream of their TSSs in mammosphere-cultured BAP1 KOs compared to MCF10A-Cas9. Down: downregulated; up: upregulated. (B) Gene ontology enrichment of downregulated genes (FDR < 0.01, log_2_FC < 0) showing ATAC loss (FDR < 0.01, log_2_FC < 0) within 2kb upstream of their TSSs in mammosphere-cultured BAP1 KOs compared to MCF10A-Cas9. Top 10 enriched ontologies are shown (p adjusted < 0.05). (C) Diamond plot of glycosylation-associated DEGs (FDR < 0.01, GO: 0070085) showing changes in chromatin accessibility within 2kb upstream of their TSSs in mammosphere-cultured BAP1 KOs compared to MCF10A-Cas9 controls. Red: ATAC gain; blue: ATAC loss. *y* axis: RNA-seq gene expression (log2FC). (D) and (E) Genome browser snapshot of ATAC-seq and H2AK119ub1 peaks in mammosphere-cultured BAP1 KOs and MCF10A-Cas9, at the *MGAT4A* and *ST6GALNAC1* genes. Horizontal bars represent differential ATAC-seq (FDR < 0.01) and/or ChIP-seq (FDR < 0.05) peaks. (F) Heatmap of glycosylation genes showing consistent alteration in chromatin accessibility (ATAC-seq, 2kb upstream of TSSs, left), RNA expression (RNA-seq, middle) and protein expression (proteomics, right) in BAP1 KO mammospheres compared to MCF10A-Cas9 controls. Colors and numbers on the heatmap represent the log2FC between BAP1 KOs and controls for each omics analysis (blue: ATAC loss or downregulation), and asterisks indicate that the change is significant (* FDR < 0.01). Where more than one ATAC-seq peak was annotated to a specific gene, only the closest peak to its TSS is shown in the heatmap.

GO analysis of downregulated genes overlapping with ATAC loss within 2kb upstream of TSSs showed enrichment in several cellular membrane-associated pathways, including top pathways previously identified by individual analysis of RNA-seq and ATAC-seq data, such as glycosylation (Figure 5B). Further assessment of chromatin accessibility and gene expression changes in genes from the glycosylation pathway (GO:0070085) revealed that differentially expressed glycosylation genes, most of them downregulated in BAP1 KOs and involved in the enzymatic addition of sugars to glycan chains (glycosyltransferases), correlate with changes in ATAC-seq signal within 2kb upstream of their TSSs (Figure 5C). Overall, the overlap between transcriptomic and ATAC-seq data suggests a correlation between reduced chromatin accessibility and downregulated gene expression in BAP1 KO mammospheres, especially in glycosylation-associated pathways.

To unravel genes and pathways that are directly regulated by BAP1’s histone deubiquitinase role, we overlapped downregulated genes, genes associated with ATAC loss (2kb upstream of TSSs), and genes associated with H2AK119ub1 gain (closest genes to H2AK119ub1 peaks) in BAP1 KO cells compared to controls. We identified 250 downregulated genes showing concomitant ATAC loss and accumulation of the silencing H2AK119ub1 mark, involved mainly in cytoskeleton and actin filament organization (Supplementary Figure S7B). The deregulation of these pathways may underpin the differences in spheroid organization and morphology displayed between controls and BAP1 KOs, and could be linked to cytoskeleton reorganization processes observed in EMT ^66^. In addition, overlap of genes associated with H2AK119ub1 gain and transcription downregulation revealed enrichment of glycosylation, cilium assembly, phospholipid, and cellular membrane pathways (Supplementary Figure S7C). These included several glycosyltransferase genes, such as *C1GALT1*, *GALNT3*, *GCNT1*, *GCNT3*, *MGAT4A* and *ST6GALNAC1*, which showed concomitant alteration in chromatin accessibility at the same regions in BAP1 KOs (Figure 5D and 5E, Supplementary Figure S8).

The impact of BAP1 loss in glycosylation-associated pathways was further confirmed at the protein level with a total proteomics approach (Supplementary Figures S9A and S9B), which revealed downregulated expression of several glycosylation proteins in BAP1 KO mammospheres (FDR < 0.01), consistent with reduced chromatin accessibility and transcription of the associated genes (Figure 5D). Moreover, total proteomics corroborated the changes in cell state and EMT, with the detected downregulation of epithelial markers EpCAM and MCAM and the upregulation of the mesenchymal marker Vimentin in BAP1 KOs, all accompanied by changes in chromatin accessibility and gene expression (Supplementary Figure S9C). Increased expression of the mesenchymal marker N-cadherin was also confirmed at the protein level upon BAP1 loss, although not linked with changes in chromatin accessibility. Finally, several genes involved in actin filament and cytoskeleton organization were found as consistently less accessible and downregulated across omics, including at the protein level (Supplementary Figure S9B). Altogether, our data reveal that loss of BAP1 leads to accumulation of H2AK119ub1 and reduced chromatin accessibility in numerous genes involved in cellular state, glycosylation and cytoskeleton organization, resulting in their downregulation at both transcript and protein levels and suggesting an impact in cell phenotypes.

### BAP1 loss disrupts glycan profiles of mammary cells

Characterization of the epigenome, transcriptome and proteome landscapes in BAP1-disrupted mammospheres revealed extensive alterations in glycosylation-associated genes, mainly involving glycosyltransferases associated with *O*-glycan and *N*-glycan biosynthesis. To examine the impact of BAP1 loss in this biological process, we adopted a glycomics approach to assess the *N*-glycan profiles of mammosphere-cultured MCF10A-Cas9 and BAP1 KO cells using hydrophilic interaction liquid chromatography (HILIC), which revealed clear changes in glycan composition between BAP1 KOs and controls. Upon BAP1 loss, we observed a significant decrease in *N*-glycan complexity, represented by higher abundance of chromatographic peaks identified at earlier retention times, while conversely, complex *N*-glycans were more abundant in MCF10A-Cas9 controls (Figure 6A and Supplementary Figure S10), pointing to an impact of BAP1 loss in the maintenance of glycan complexity in mammary cells.

**Figure 6.**
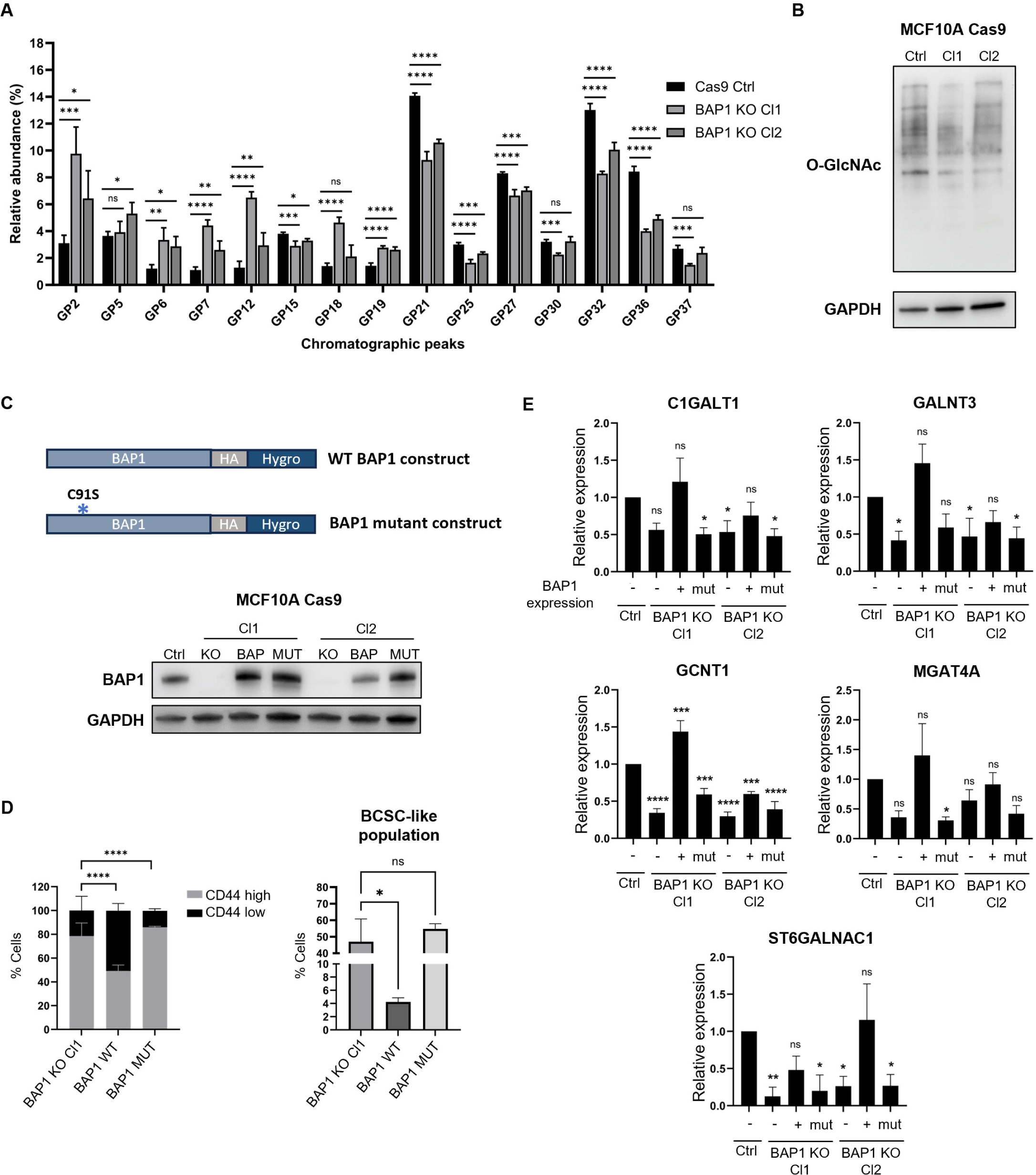
BAP1 loss disrupts glycan profiles, and its rescued activity restores phenotypes. (A) Relative abundance of chromatographic peaks identified by *N*-glycan profiling of mammosphere-cultured MCF10A-Cas9 and BAP1 KO cells. Significance analysis performed by Dunnett’s test (n=4, * *P* < 0.05, ** *P* < 0.01, *** *P* < 0.001, **** *P* < 0.0001, ns: not significant). Top most abundant peaks across samples are shown. GP: glycan peak. (B) *O*-linked *N*-acetylglucosamine (O-GlcNAc) expression levels in MCF10A-Cas9 controls (Ctrl) and two BAP1 knockout (KO) clones (Cl1 and Cl2) cultured as mammospheres. GAPDH levels were assessed as an internal control. (C) Top: schematic representation of the constructs used for BAP1 re-expression, containing an HA tag and hygromycin (Hygro) resistance. Representations of the wildtype (WT) and mutant BAP1 (C91S mutation) constructs are shown. Bottom: BAP1 protein expression levels in MCF10A-Cas9 controls (Ctrl), and two BAP1 knockout (KO) clones (Cl1 and Cl2) infected with BAP1 wildtype (BAP) and BAP1 C91S mutant (MUT) rescue plasmids. GAPDH levels were assessed as an internal control. (D) Left: distribution of CD44 high and low populations in BAP1 KO Cl1 and BAP1 rescue cells (wildtype: WT; and mutant: MUT) cultured as mammospheres. Significance analysis performed by Fisher’s exact test on absolute cell counts (n=2, **** *P* < 0.0001). Right: proportion of BCSC-like populations based on expression of CD44, CD24 and EpCAM in BAP1 KO Cl1 and BAP1 rescue cells (wildtype: WT; and mutant: MUT) cultured as mammospheres. Significance analysis performed by Dunnett’s test (n=2, * *P* < 0.05, ns: not significant). (E) Expression of glycosyltransferase genes *C1GALT1*, *GALNT3*, *GCNT1*, *MGAT4A* and *ST6GALNAC1* in MCF10A-Cas9 (Ctrl), BAP1 KO cells and BAP1 rescues (wildtype, +; and mutant, mut) cultured as mammospheres. Significance analysis performed by Dunnett’s test compared to control (Ctrl) sample (n=3, * *P* < 0.05, ** *P* < 0.01, *** *P* < 0.001, **** *P* < 0.0001, ns: not significant).

Because BAP1 may interact with OGT, a glycosyltransferase which catalyzes the addition of *O*-linked *N*-acetylglucosamine (*O*-GlcNAc) to proteins ^67^, we assessed *O*-GlcNAc levels in BAP1 KOs and controls. Even though BAP1 loss did not alter the accessibility and expression of the *OGT* gene, BAP1 KOs exhibited reduced levels of *O*-GlcNAc compared to control mammospheres (Figure 6B), hinting to an impact of BAP1 deregulation in *O*-glycosylation levels. Together, these results indicate that chromatin accessibility changes resulting from BAP1 loss and the associated reduction in the expression of glycosylation genes and proteins impacts both glycan profiles and glycan abundance at a global level in non-tumorigenic mammary cells.

### Rescued expression of BAP1 restores BCSC marker and glycosylation gene levels in a catalytic activity-dependent manner

To confirm if changes in mammary cell phenotype and glycosyltransferase gene expression are linked to BAP1’s deubiquitinase role, we restored BAP1 expression in both KO clones using either a wildtype *BAP1* construct, or *BAP1* with an inactivating mutation in its ubiquitin carboxy hydrolase domain (C91S) ^68^ (Figure 6C). Restoration of wildtype BAP1 led to re-established surface expression levels of the BCSC marker CD44 in mammospheres, with a decrease in the CD44^high^ population from over 75% to around 50% of cells, while the absence of catalytic activity failed to rescue normal CD44 levels (Figure 6D). The proportion of mesenchymal BCSC-like cells was also markedly reduced upon BAP1 restoration, but remained at similar levels to BAP1 KOs in the presence of mutant BAP1 (Figure 6D), suggesting that enhanced expression of BCSC markers upon BAP1 loss is dependent on its deubiquitinase activity.

In addition, we demonstrated that the expression of several glycosyltransferase genes is reduced upon BAP1 loss in non-tumorigenic mammospheres, and that this reduction is dependent on BAP1’s catalytic activity (Figure 6E). Of note, BAP1 rescue also led to increased or re-established levels of different cell cycle genes which were identified as downregulated upon BAP1 loss (Supplementary Figure S11). Still, the expression of a few glycosylation and cell cycle-associated genes, such as *GCNT1* and *EREG*, was only partially rescued in the presence of wildtype BAP1, suggesting that expression of these genes may also be co-regulated by other mechanisms. Overall, these results show that normal catalytic activity of BAP1 is needed to maintain adequate expression levels of CD44, EpCAM and CD24 markers, as well as cell cycle genes and several glycosyltransferases.

## DISCUSSION

In this study, we applied a comprehensive *in-vitro* CRISPR/Cas9 screening approach in non-tumorigenic breast cells to identify potential epidrivers of the acquisition of BCSC markers, a phenotype associated with BC cell plasticity. Our approach led to the identification of BAP1 deregulation as a putative epidriver of BCSCs. Extensive validation and multi-omics characterization in mammary spheroids confirmed our observation and further revealed that BAP1 loss impacts cell morphology and phenotype and leads to broad epigenome, transcriptome, and proteome deregulation. Additionally, we linked the deubiquitinating activity of BAP1 to the control of glycosylation-associated gene expression and revealed glycomic aberrations in mammary cells as a result of BAP1 loss.

This is the first report to unveil *BAP1* disruption as an epigenetic driver of mammary cell plasticity and the emergence of mammary cells with mesenchymal BCSC-like markers (CD44^high^ CD24^low^ EpCAM^-^) and altered expression of EMT-associated genes. This includes the reduced expression of epithelial markers EpCAM and MCAM, linked to changes in chromatin accessibility, and the upregulation of mesenchymal markers such as *ZEB1*, a transcription factor that induces cell plasticity through transcriptional repression of tumor suppressor genes and activation of EMT ^69^. Moreover, our observation that *BAP1* is a frequently mutated ERG in TNBCs, a BC subtype enriched in BCSCs ^55,56^, and that its reduced expression in BC tumors associates with lower disease-free survival, suggest that *BAP1* expression may be linked to less aggressive phenotypes in BCs. Indeed, in different cancer types, BAP1 disruption and loss have been suggested to contribute to aggressiveness ^70^, but the effects of its disruption in BC are still largely unexplored. Here, our analyses have shown that BAP1 loss alone is sufficient for the acquisition of mammary cells expressing BCSC markers, and revealed several other putative epidrivers of this phenotype, all involved in histone modification. Remarkably, three of these putative epidrivers (*BAP1*, *ASXL2* and *KDM6A*) counteract gene silencing promoted by Polycomb-group complexes ^54,63,71^, suggesting that disruptions in this epigenetic mechanism may be implicated in mammary cell plasticity and the BCSC phenotype.

BAP1 deletion in mammary cells unveiled extensive deregulation in their epigenetic profile, mainly associated with loss in chromatin accessibility, consistent with BAP1’s role as a histone modifier and transcriptional activator, and in line with previous reports of chromatin condensation and reduced accessibility upon BAP1 loss in other cell types ^72-74^. Furthermore, we showed that increased H2AK119 ubiquitination levels following BAP1 loss largely overlapped with less accessible chromatin surrounding TSSs and transcriptional downregulation, revealing novel chromatin targets for BAP1 in mammary cells, including cell cycle, glycosylation, and cellular organization-associated genes. Chromatin accessibility changes observed following BAP1 loss also revealed a possible impact in the binding of different transcription factors to chromatin. This included overrepresentation of TP73 and TP53 motifs in BAP1 KOs, involved in regulation of stemness, self-renewal and cellular differentiation ^75,76^, and TWIST1, a well-known inducer of EMT and stemness in tumor initiation and progression ^77^. Interestingly, top underrepresented motifs in BAP1 KOs belong to the FOX family of transcription factors, involved in various functions such as regulation of cell cycle, metabolism, and stem cell maintenance ^78^. One of the FOX transcription factors identified in our analysis, FOXK2, may recruit BAP1 to its target genes for the regulation of their expression through BAP1’s histone deubiquitinase function ^64,79^, and was shown to inhibit the proliferative and invasive capacities of BC cells, suggesting a tumor suppressive role ^80^. Here, the finding of extensive deregulation of transcription factor binding potential upon BAP1 loss, with increased binding of stemness-promoting factors and decreased binding of transcription factors with tumor suppressive roles, may be underpinning the shift in cell identity and the stemness-associated phenotypes observed in BAP1 KOs.

Our results showing an accumulation of BAP1 KOs in the S phase of the cell cycle are consistent with previous reports linking BAP1 disruption to delayed cell cycle progression ^61,65,67,81,82^. In addition, reduced proliferation is a common characteristic of mesenchymal BCSCs ^13^, further implying that BAP1 loss may lead to the emergence of cells with this phenotype. Mechanistically, BAP1 may promote cell cycle progression and proliferation by H2AK119 deubiquitination of E2F target genes, resulting in their activation and transcription ^57,58^. Moreover, partial or full restoration of cell division-associated gene expression upon rescue of catalytically active BAP1 further implicates its histone deubiquitinase function as a key mechanism in normal cell cycle regulation in mammary cells.

Another striking finding from our in-depth characterization of BAP1 disruption in mammospheres was the overall deregulation of glycosylation processes, observed at the epigenome, transcriptome, proteome, and phenotypic levels. Previous studies reported that BAP1 mutations may impair its interaction with OGT and result in reduced levels of *O*-GlcNAc in mesothelioma and uveal melanoma ^83^, and that mutant BAP1 tumors associate with a specific glycan signature in renal cell carcinoma ^84^. Furthermore, we observed that BAP1-deficient tumors (BCs and pan cancer) show lower expression levels of several glycosylation pathway genes when compared to tumors exhibiting normal or low-level gain in BAP1 copy number (Supplementary Figure S12). This further supports an association between BAP1 function and the regulation of glycosylation that warrants further exploration. To our knowledge, the regulation of cellular glycosylation and glycan levels and complexity by BAP1 through epigenetic mechanisms had not been previously described, notably in mammary cells. Here, we reveal that changes in glycosylation pathways and glycan levels observed following BAP1 loss are largely linked to epigenetic and chromatin remodeling. In addition, we show rescued expression of several glycosyltransferases upon re-established BAP1 activity, further strengthening the notion that BAP1 maintains normal glycosylation levels through the epigenetic regulation of glycosylation-associated genes.

Glycans are complex carbohydrate chains that can be attached to different molecules, with roles in regulation of cell metabolism, proliferation, cell adhesion, and cell signaling ^85,86^. In cancers and cancer stem cells, aberrant glycosylation is commonly found either in the form of truncated glycosylation in early cancer development, characterized by impaired synthesis of complex glycans, or as increased glycan branching in cancer progression ^87^, contributing to aggressive phenotypes, tumor progression, metastasis, and therapeutic resistance ^86,88^. Different cancer stem cell markers, such as CD44, are extensively modified by glycosylation, and alterations in their glycan profiles may impact their stability, ligand binding and cell signaling activation ^88^. The concomitant deregulation of glycosylation and the emergence of cells with BCSC markers upon BAP1 loss suggest a possible mechanistic link between these two events. However, further studies are warranted to better characterize the nature of altered glycan structures upon BAP1 loss, its impact on the expression of BCSC markers and its effect on the functions of BCSCs.

Our observations place BAP1’s epigenetic activity and H2AK119ub1 removal as important aspects in the maintenance of mammary cell identity. In different cancer types, increased activity of Polycomb complex PRC1, which catalyzes H2AK119ub1, is associated with oncogenic roles, malignant transformation and stemness ^89-92^. Our results suggest that similar to PRC1 overactivation, BAP1 loss may also contribute to cell plasticity and stemness-associated phenotypes, possibly through the accumulation of H2AK119 ubiquitination, with an impact on cell cycle and glycosylation pathways. Thus, a proper balance between BAP1 and PRC1 activity in regulating H2A ubiquitination levels may play a major role in preserving mammary cell state and in limiting a shift towards a possible oncogenic fate.

## CONCLUSIONS

Our results reveal novel mechanistic roles for BAP1 as an epidriver of BCSC marker acquisition, which may be further explored for the understanding of cancer stem cell biology, BC initiation, and for the assessment of BAP1 disruption in aggressive breast cancers, such as TNBCs. This study may open novel avenues for exploring the effect of epigenetic drugs in the control of H2AK119 ubiquitination levels, including PRC1 inhibitors ^93^, as a potential therapeutic approach to eliminate cells with cancer stem cell characteristics in breast cancers.

## LIST OF ABBREVIATIONS

2D/3D: two-dimensional/three-dimensional
ATAC: assay for transposase-accessible chromatin
bp: base pairs
BC: breast cancer
BCSC: breast cancer stem cell
ChIP: chromatin immunoprecipitation
Cl: clone
CRISPR: clustered regularly interspaced short palindromic repeats
CSC: cancer stem cell
Ctrl: control
DEG: differentially expressed gene
ECAR: extracellular acidification rate
EMT: epithelial-to-mesenchymal transition
ERG: epigenetic regulator gene
FC: fold change
FDR: false discovery rate
GO: gene ontology
gRNA: guide RNA
GSEA: gene set enrichment analysis
H&E: hematoxylin and eosin
Kb: kilobases
KO: knockout
LOF: loss-of-function
MET: mesenchymal-to-epithelial transition
MOI: multiplicity of infection
OCR: oxygen consumption rate
PCA: principal component analysis
PR-DUB: Polycomb repressive deubiquitinase
PRC1/2: Polycomb repressive complex 1 or 2
RT-qPCR: quantitative reverse transcription polymerase chain reaction
TNBC: triple negative breast cancer
TSS: transcription start site
WT: wildtype

## DECLARATIONS

### Ethics approval and consent to participate

Not applicable.

### Consent for publication

Not applicable.

### Data availability

High-throughput sequencing data for the CRISPR screen, ATAC-seq, ChIP-seq and RNA-seq experiments are available from Gene Expression Omnibus (GEO) (accession numbers GSE271680, GSE271681, GSE271682, GSE271683, GSE271684). Mass spectrometry proteomics data have been deposited to the ProteomeXchange Consortium via the PRIDE repository with the dataset identifier PXD057440.

## Acknowledgements

The authors would like to thank Drs. Anne Pierre-Morel, Maria Ouzounova and Rahma Benhassoun for their scientific and technical support; the Cancer Research Centre of Lyon (CRCL) flow cytometry facility for the support with cell sorting and flow cytometry experiments; the members of the Quantitative Proteomics Unit at IBC2 (Goethe University, Frankfurt), in particular Martin Adrian-Allgood and Bhavesh Parmar for technical help and Kristina Wagner for preparing LC columns; Dr. Ivan Đikić for the scientific support with proteomics experiments; and Dr. Mirjam Zeisel for kindly providing the anti-O-GlcNAc antibody. We thank Elizabeth Page and Foster Jacobs for kindly proofreading the manuscript.

## Disclaimer

Where authors are identified as personnel of the International Agency for Research on Cancer/World Health Organization, the authors alone are responsible for the views expressed in this article and they do not necessarily represent the decisions, policy, or views of the International Agency for Research on Cancer/World Health Organization.

## Authors’ contributions

M.G.D.S.A. performed experiments and bioinformatic analyses, analyzed data, and wrote the manuscript; A.S. and C.Cu. performed experiments; V.C., C.R., C.P.P. and S.K. performed bioinformatic analyses of transcriptomic and epigenomic data; T.M. performed proteomic analysis; C.Ca. generated and processed histological sections; G.G.L. analyzed histological sections; L.P. generated Cas9-expressing cell lines; E.B-F., S.R. and S.K. performed survival analysis; A.G. provided support with statistical analyses; S.H., M.P-B. and G.L. performed glycan profiling; E.C. performed metabolic assays; Z.H. designed and supervised the study, and contributed to manuscript writing; R.K. designed and supervised the study, performed experiments, analyzed data, and contributed to manuscript writing; all authors reviewed the manuscript.

