## Supplementary information for "Disruption of the epigenetic regulator BAP1 drives chromatin remodeling leading to the emergence of cells with breast cancer stem cell properties and aberrant glycosylation"

**This document contains Supplementary Methods, Supplementary Figures S1-S12, and Supplementary Tables S1-S8.**

#### SUPPLEMENTARY METHODS

##### Cell and mammosphere culture

MCF10A cells (ATCC CRL-10317) were maintained in DMEM/F12 medium supplemented with 5% fetal bovine serum, insulin (10 µg/mL), epidermal growth factor (20 ng/mL), hydrocortisone (500 ng/mL), cholera toxin (100ng/mL), and 1% penicillin/streptomycin. Cell culture was performed at 37°C in a humidified, 5% $CO_2$ atmosphere.

Cells were also maintained in culture as mammospheres (mammary spheroids), a three-dimensional (3D) low-attachment culture condition which allows enrichment of breast (cancer) stem cells [1,2]. For this, MCF10A cells were seeded at a density of $1x10^4$ cells/$cm^2$ in 100mm dishes coated with 20 mg/mL Poly(2-hydroxyethyl methacrylate) (Sigma-Aldrich), and cultured in DMEM/F12 medium as described above. Single size-normalized mammospheres were generated by seeding 3 - $5x10^4$ cells in 96-well round bottom ultra-low attachment plates (Corning), followed by 3-5 minutes centrifugation at 1,000 rpm to pull cells to the bottom of wells. Mammospheres were kept in culture for seven days unless stated otherwise. For the generation of single cell suspensions, mammospheres were dissociated with 0.25% Trypsin EDTA (Thermo Fisher Scientific) and gentle mixing by pipetting.

##### Western Blot

Protein expression was assessed using 20-30 µg of total protein extracted with RIPA buffer and protease inhibitors (cOmplete Protease Inhibitor Cocktail, Roche). Samples were loaded in precast polyacrylamide gels (Bio-Rad) and transferred to polyvinylidene fluoride (PVDF) membranes (Bio-Rad). Membranes were incubated overnight at 4°C with antibodies for Cas9 (1:2,000, ab191468, abcam), BAP1 (1:500, sc-28383, Santa Cruz Biotechnology), O-GlcNAc (1:1,000, MA1-072, Invitrogen) and GAPDH (1:2,000, sc-32233, Santa Cruz Biotechnology) as a loading control, followed by incubation with appropriate species-matched secondary antibodies linked to HRP (Agilent Dako) and detection with Clarity Western ECL mix (Bio-Rad). For the analysis of H2AK119ub1 levels, histones were precipitated using the Histone Extraction Kit (Active Motif) according to the manufacturer's instructions, and western blots were performed as described above, using Ubiquityl-Histone H2A (Lys119) antibody (1:2,000,

D27C4, Cell Signaling Technology) and anti-histone H3 antibody (1:2,000, ab1791, abcam) as a loading control.

##### Histological evaluation and imaging of mammospheres

Histological analysis was performed in size-normalized mammospheres cultured for 4-5 days (initial plating concentration of $3x10^4$ cells per mammosphere). Mammospheres were collected, washed with 1X PBS, and fixed with 4% formaldehyde solution for 45 minutes, followed by two washes with 1X PBS. Paraffin embedding and sectioning of spheroids was performed following a previously described protocol [3], and 3 µm sections were processed for hematoxylin and eosin (H&E) staining. Slides were scanned with the Glissando Slide Scanner (Objective Imaging) using 40× magnification and examined by an experienced pathologist (IARC's histopathology platform).

##### Validation of cell surface marker expression

Expression of cell surface proteins was analyzed through flow cytometry (n=3) using the CD44 (1:400 dilution), CD24 (1:100 dilution) and EpCAM (1:200 dilution) antibodies previously employed in cell sorting. MCF10A-Cas9, BAP1 KO and BAP1 rescue cells were kept in culture as mammospheres for seven days and trypsinized to obtain single cells. A density of $3 – 4x10^5$ cells per staining was incubated in 100 µL of antibody staining solution (PBS 2% FBS with antibodies) for a minimum of 1 hour at 4°C, protected from light. After antibody incubation, cells were washed three times with PBS 2% FBS and analyzed in a flow cytometer (BD LSRFortessa). Cells were also stained with isotype control antibodies (1:400, APC 130-113-434, FITC 130-113-437, PE 130-113-438, Miltenyi Biotech) for the compensation of fluorescence signals. Quantification of cell populations based on fluorescence signals was performed with FlowJo v10 software. Fisher's exact test was used to assess for differences in CD44-high and -low expression groups, and one-way ANOVA with Dunnett's multiple comparisons test was used to assess differences in representation of specific populations (p-value threshold < 0.05).

##### Quantitative RT-PCR (RT-qPCR)

Total RNA was extracted using either AllPrep DNA/RNA Mini Kit or RNeasy Mini Kit (Qiagen). 500 ng - 1 µg of total RNA were used for reverse transcription with SuperScript III First-Strand Synthesis Super Mix (Invitrogen). RT-qPCR was performed with SsoAdvanced Universal SYBR

Green Supermix (Bio-Rad) in CFX Real-Time PCR Detection Systems (Bio-Rad), and *GAPDH* expression was used as a housekeeping gene control for normalization of results. Relative expression was calculated using the ΔΔCt method. Primers used in RT-qPCR analyses are listed in Supplementary Table S2. Three independent RT-qPCR experiments were performed for each gene assessed (n=3), and statistical analysis was performed using one-way ANOVA with Dunnett's multiple comparisons test, with a p-value threshold of $< 0.05$.

**Pyrosequencing**

Total DNA extracted with AllPrep DNA/RNA Mini Kit (Qiagen) was quantified with Qubit (Thermo Fisher Scientific), and 500 ng of DNA were bisulfite-converted with the EZ DNA Methylation kit (Zymo Research) following the manufacturer's protocol. Target regions were amplified by PCR and pyrosequenced using PyroMark Gold reagents on a PyroMark Q96ID instrument (Qiagen). The list of primers used for pyrosequencing is available from Supplementary Table S3. Pyrosequencing was performed in duplicates (n=2), and percentage of methylation for each CpG position was calculated as the mean methylation of the replicates. To evaluate differential methylation in the whole region, linear regression analysis (R v4.1.2) was performed for each CpG and comparison made, followed by meta-analysis with METAL (Meta Analysis Helper) software [4], with a p-value threshold of $< 0.05$.

**Proliferation assays**

Assessment of proliferation in attached cells was performed with the CellTiter 96 Aqueous One Solution Cell Proliferation Assay (Promega), as per the manufacturer's instructions. Briefly, MCF10A control and BAP1 KO cells were plated in 96-well plates at a density of $1x10^3$ cells/well, in 100 µL of culture medium. At each timepoint (24h, 48h, 72h), 20 µL of CellTiter reagent was added to each well, and, after 3h of incubation at 37°C in a 5% $CO_2$ atmosphere, absorbance was measured at 492nm. Wells containing culture medium and CellTiter reagents were used as blanks for absorbance measurements. Proliferation assays were performed in triplicates (n=3), and differences between samples were assessed using two-way ANOVA with Dunnett's multiple comparisons test, with a p-value threshold of $< 0.05$.

**Cell cycle analysis**

Cell cycle analysis was performed by propidium iodide (PI) staining of mammosphere-cultured cells in triplicates (n=3). Briefly, MCF10A-Cas9 and BAP1 KO cells were kept in culture as mammospheres for seven days and trypsinized to obtain single cells. Single cell suspensions were fixed with 70% EtOH for 1h at 4°C, then incubated with RNAse A (100 µg/mL) and PI (40 µg/mL) in PBS for 30 minutes at 37°C, protected from light. After incubation, cells were analyzed in a flow cytometer (BD LSRFortessa). Quantification of cell cycle phases was performed using the Dean-Jett-Fox model on FlowJo v10 software. One-way ANOVA with Dunnett's multiple comparisons test was used to calculate differences in representation of each cell cycle phase between samples, with a p-value threshold of $< 0.05$.

**Metabolic assays**

Assessment of metabolic changes was performed in MCF10A-Cas9 and MCF10A-Cas9-BAP1 KO cells cultured as single, size-normalized mammospheres, at a density of $5x10^4$ cells/spheroid. Briefly, mammospheres were kept in culture for 7 days, transferred to poly-L-lysine-coated (1:10, coated for 1 hour at 37°C) Seahorse XFe96 Spheroid Microplates (Agilent), and analyzed using Seahorse XF Cell Mito Stress Test and Glycolysis Stress Test Kits (Agilent) as per manufacturer's protocol. Oxygen consumption rate (OCR) and extracellular acidification rate (ECAR) measurements were normalized to protein expression of each spheroid using Pierce BCA Protein Assay Kit (Thermo Fisher Scientific). Metabolic assays were performed in four independent replicates (n=4) for the Glycolysis Stress Test and eight independent replicates (n=8) for the Mito Stress Test. Each replicate represents the mean of measurements of at least four individual spheroids per cell type.

**Integrative analyses**

Integration of transcriptomic and epigenomic data was performed with differentially expressed genes and proteins from RNA-seq and proteomics analyses (FDR < 0.01), differentially accessible ATAC-seq peaks (FDR < 0.01), and differential H2AK119ub1 peaks (FDR < 0.05) following BAP1 loss in MCF10A cells. Differential ATAC-seq and ChIP-seq peaks were annotated to the closest gene for the Venn overlaps and pathway enrichment analyses.

Diamond plots of ATAC-seq and RNA-seq were generated as previously described [5]. Briefly, each diamond represents a differential ATAC-seq peak annotated to a specific gene, and diamond color associates with direction of chromatin accessibility change in BAP1 KOs (red for gain in accessibility and blue for loss). Position of the bottom-most diamond in relation to the *y* axis represents the $\log_2$FC (FC, fold change) in expression of the associated gene.

Heatmaps were prepared with the package "pheatmap" v1.0.12 [6] in R (v4.1.2). For RNA-seq expression heatmaps (Supplementary Figure S6), normalized RNA-seq read counts obtained from DESeq2 were log10-transformed, and z-scores were calculated for each gene across samples. Heatmap colors represent the Z-score obtained for each gene in each sample. Hierarchical clustering of samples (columns) was performed using Euclidean distance. For heatmaps of changes in chromatin accessibility (ATAC-seq, 2kb upstream of TSS), RNA expression (RNA-seq) and protein expression (proteomics) in specific genes (Figure 5F, Supplementary Figure S9C), log2FC and FDR values obtained from differential analyses between BAP1 KO mammospheres and MCF10A-Cas9 controls were used. Where more than one ATAC-seq peak annotated to a same gene of interest, only the closest peak to the TSS was considered. Heatmap colors and numbers represent the log2FC values, and asterisks (*) indicate whether the observed change was considered as significant (FDR < 0.01).

**SUPPLEMENTARY FIGURES**

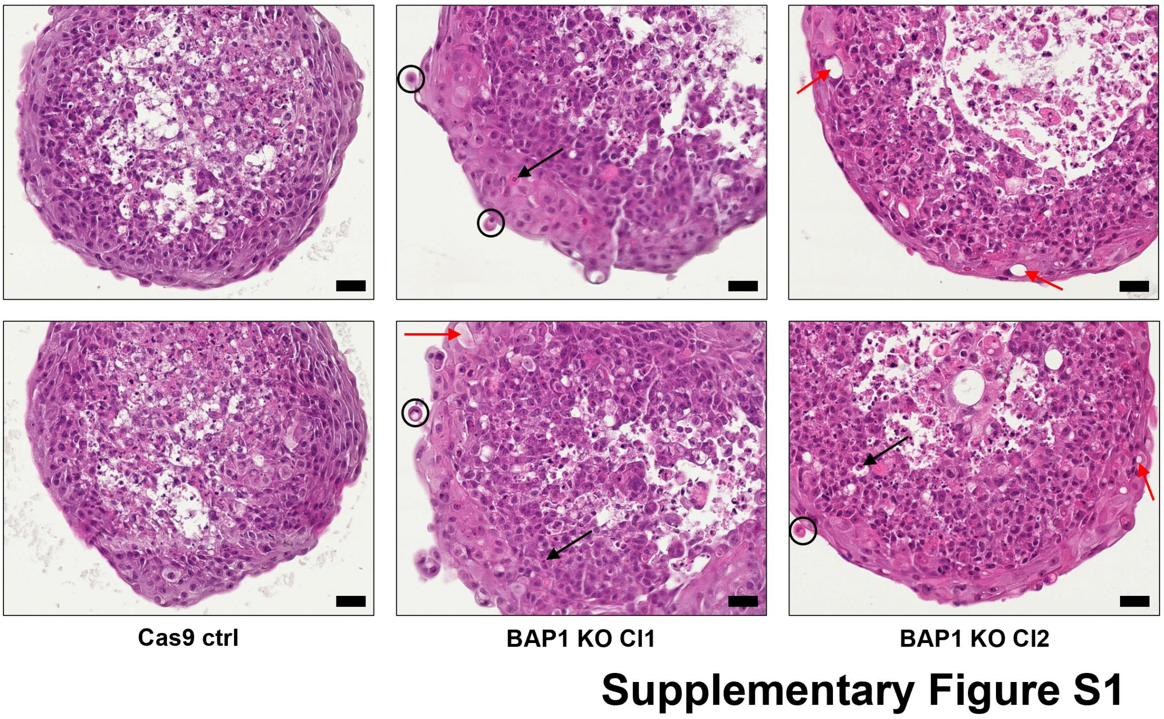


**Supplementary Figure S1.** Representative images at 30x magnification of single, size-normalized MCF10A-Cas9 control and BAP1 KO mammospheres stained with hematoxylin and eosin show detailed intracytoplasmic vacuolization (red arrows), apoptotic bodies (black arrows) and cellular decohesion (black circles), noted mainly at the periphery of the mammospheres. Scale bars: 33 µm.

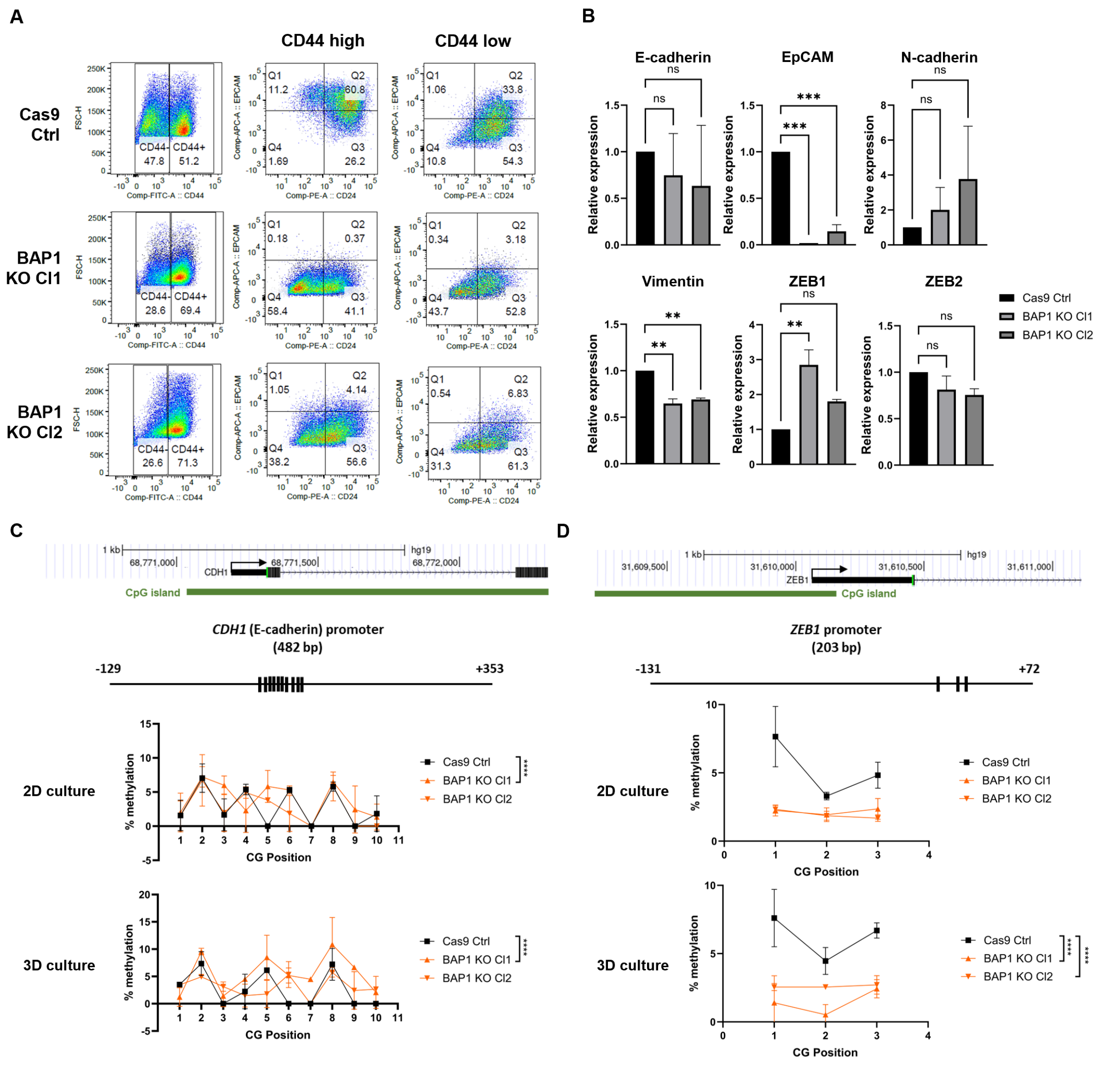


#### Supplementary Figure S2

**Supplementary Figure S2.** (A) Validation of the acquisition of mesenchymal BCSC-like markers in MCF10A-Cas9 BAP1 KO cells cultured as mammospheres by flow cytometry staining. Samples were stained for CD44, CD24 and EpCAM markers. Representative image of one replicate. (B) Expression of EMT-associated genes in MCF10A-Cas9 and BAP1 KO cells cultured in attachment. Significance analysis performed by Dunnett's test (n=2, ** $P$ < 0.01, *** $P$ < 0.001, ns: not significant). (C-D) Pyrosequencing analysis at *CDH1* (E-cadherin) (C) and *ZEB1* (D) promoters in BAP1 KOs and MCF10A-Cas9 cells cultured in attachment (2D culture) and as mammospheres (3D culture). Top: coordinates of promoter location of *CDH1* and *ZEB1* in the human genome, CpG island regions shown in green. Middle: relative positions of CpGs in the amplicons. Bottom: representation of methylation levels at each CpG locus. Significance analysis performed by linear regression analysis for each CpG and comparison made, followed by meta-analysis with METAL software (n=2, **** $P$ < 0.0001).

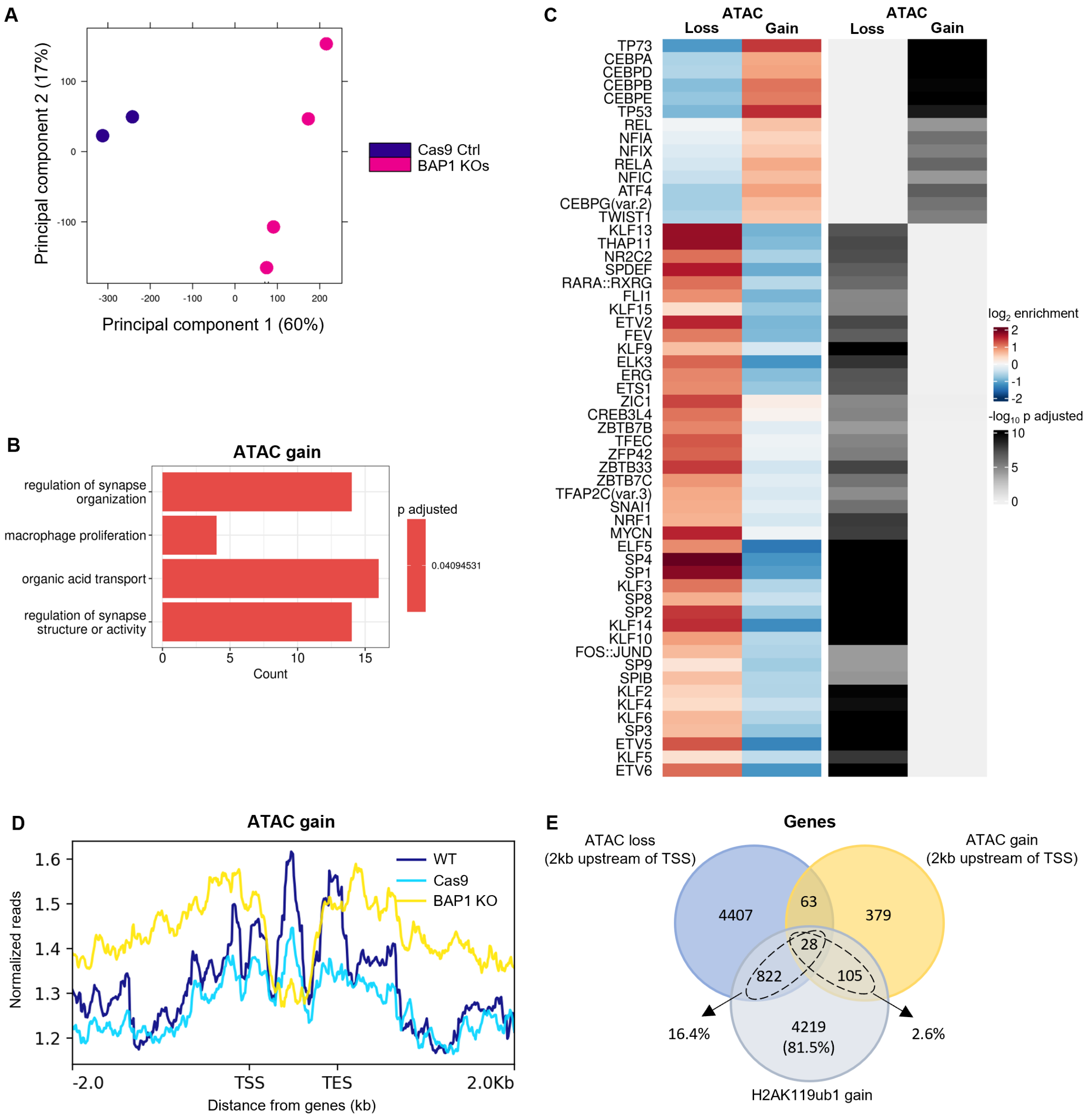


### Supplementary Figure S3

**Supplementary Figure S3.** (A) Principal component analysis (PCA) of ATAC-seq samples of BAP1 KOs and MCF10A-Cas9 cells cultured as mammospheres (n=2, two BAP1 KO clones analyzed). (B) Gene ontology enrichment of genes associated with differential ATAC gain peaks (FDR < 0.01, log2FC > 0) falling 2kb upstream from TSSs in BAP1 KOs compared to MCF10A-Cas9 cells cultured as mammospheres. All enriched gene ontologies are shown (p adjusted < 0.05). (C) Transcription factor binding enrichment on differential ATAC-seq peaks (FDR < 0.01) falling 2kb upstream from TSSs in BAP1 KOs compared to MCF10A-Cas9 cells cultured as mammospheres. Top 56 transcription factor motifs are shown (p adjusted < 0.05). (D) Distribution of H2AK119ub1 ChIP-seq peaks (FDR < 0.05) in regions of increased ATAC-seq accessibility (ATAC gain, FDR < 0.01, log2FC > 0) identified in BAP1 KOs compared to MCF10A-Cas9 cells. Regions within 2kb upstream and 2kb downstream of TSSs and transcription end sites (TESs), respectively, are shown. ChIP-seq reads were normalized with RPGC per bin (reads per genome coverage per bin) normalization method. Distribution of ChIP-seq peaks was analyzed for MCF10A wildtype cells (MCF10A-WT), MCF10A-Cas9 control cells (MCF10A-Cas9), and both BAP1 KO clones (BAP1 KO). ATAC-seq was performed in mammosphere-cultured cells, and ChIP-seq in attached cells (n=2). (E) Venn diagram of overlap between genes associated with H2AK119ub1 gain (FDR < 0.05) and genes associated with ATAC loss and/or ATAC gain (FDR < 0.01) within 2kb upstream of their TSSs in BAP1 KOs compared to MCF10A-Cas9 control cells.

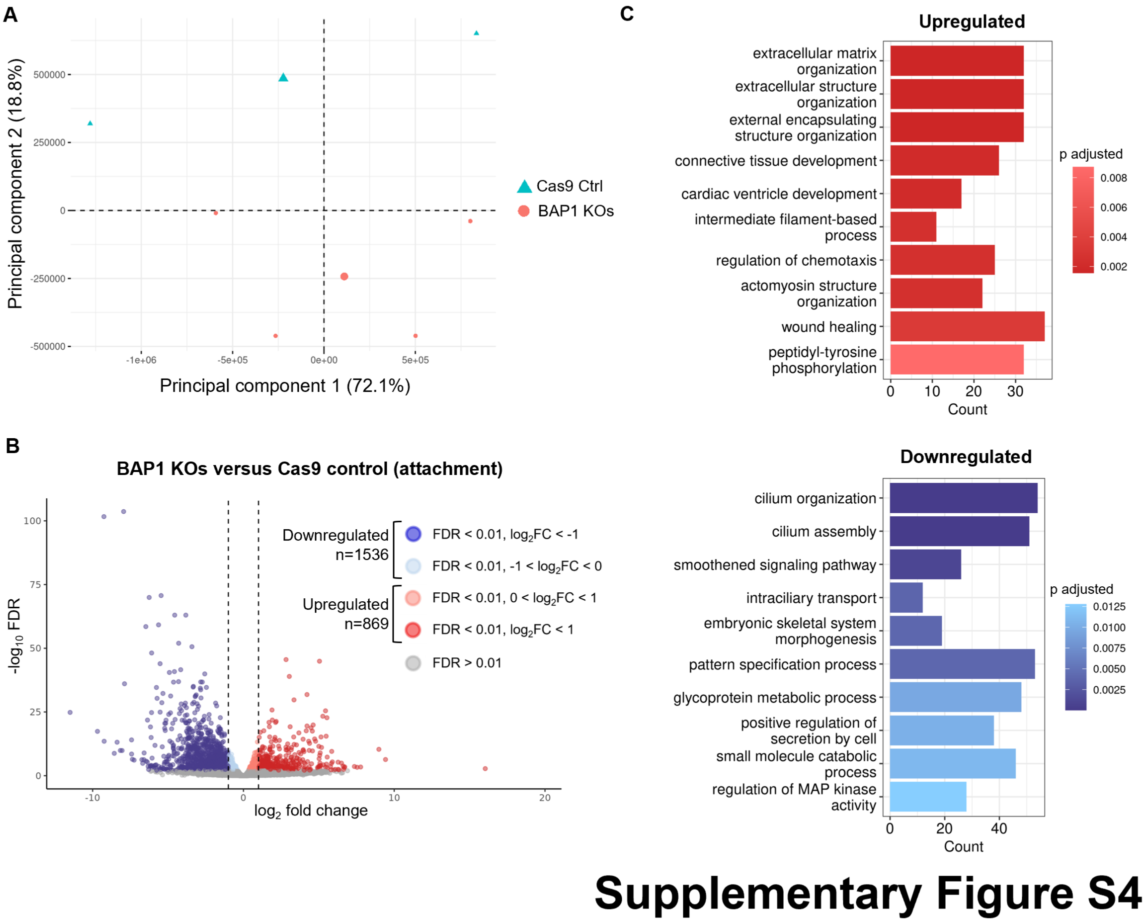


**Supplementary Figure S4.** (A) PCA of RNA-seq samples of BAP1 KOs and MCF10A-Cas9 cells cultured as mammospheres (n=2, two BAP1 KO clones analyzed). (B) Volcano plot of differential expression analysis in BAP1 KOs compared to MCF10A-Cas9 control cells cultured in attachment (n=2 for MCF10A-Cas9 cells and n=4 for BAP1 KOs). Blue and red-colored dots represent differentially expressed genes (DEGs, FDR < 0.01) with reduced or increased expression (log2FC < 0 and log2FC > 0), respectively. (C) Gene ontology enrichment of upregulated (top) and downregulated (bottom) DEGs (FDR < 0.01) in BAP1 KOs compared to MCF10A-Cas9 cells cultured in attachment. Top 10 enriched gene ontologies are shown (p adjusted < 0.05).

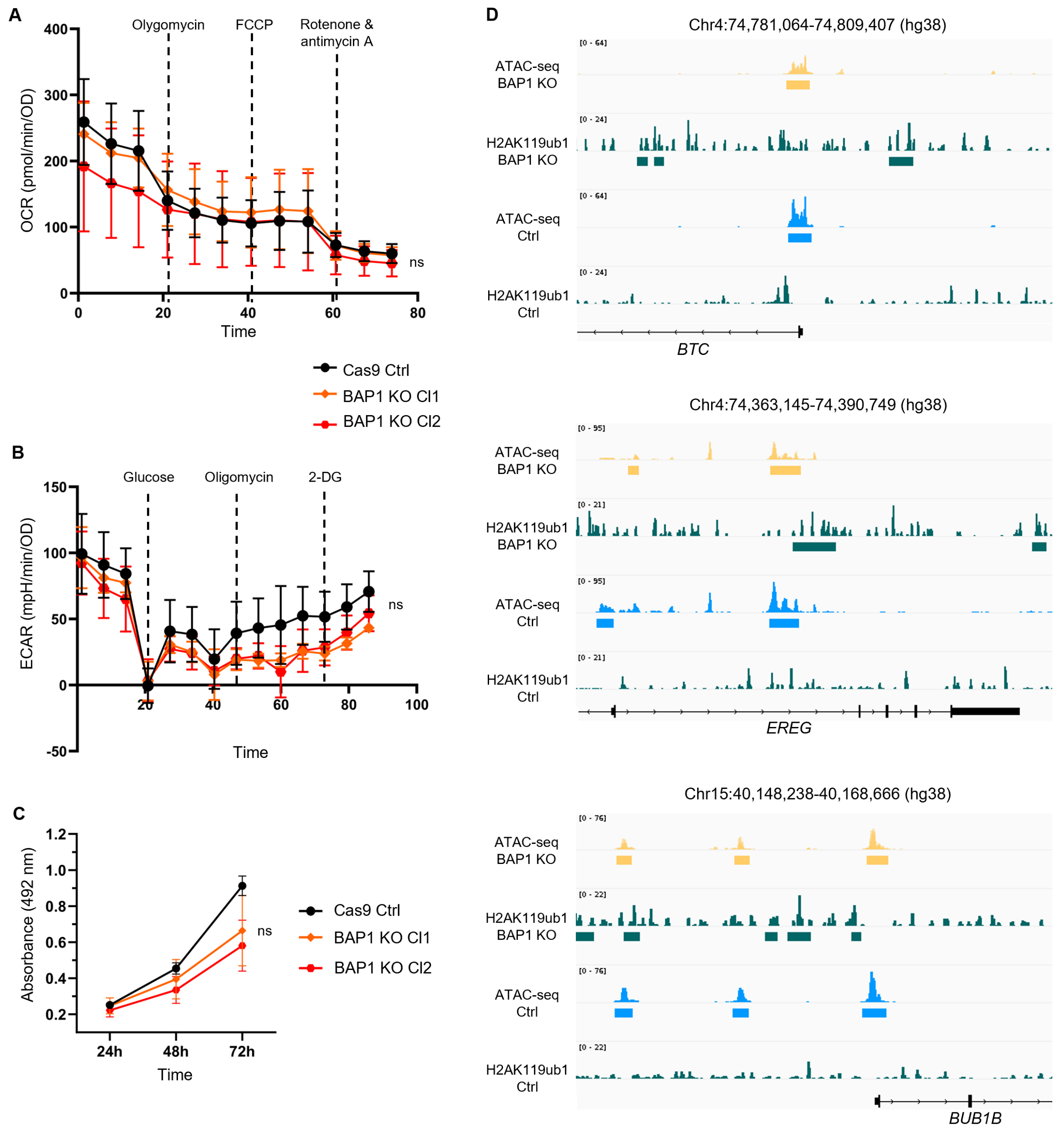


### Supplementary Figure S5

**Supplementary Figure S5.** (A) Assessment of mitochondrial function by oxygen consumption rate (OCR) measurement in BAP1 KOs and MCF10A-Cas9 mammospheres. Measurements normalized to protein expression levels of each spheroid. Significance analysis performed by Dunnett's test (n=8, ns: not significant). (B) Assessment of glycolytic capacity by extracellular acidification rate (ECAR) measurement in BAP1 KOs and MCF10A-Cas9 mammospheres. Measurements normalized to protein expression levels of each spheroid. Significance analysis performed by Dunnett's test (n=4, ns: not significant). (C) Proliferation curves of BAP1 KOs and MCF10A-Cas9 cells cultured in attachment for 72h, assessed with MTS assay. Significance analysis performed by Dunnett's test (n=3, ns: not significant). (D) Genome browser snapshot of ATAC-seq and H2AK119ub1 peaks in BAP1 KOs and MCF10A-Cas9 controls cultured as mammospheres, at the *BTC*, *EREG* and *BUB1B* genes. Colored horizontal bars represent differential peaks identified in ATAC-seq (FDR < 0.01) and/or ChIP-seq (FDR < 0.05).

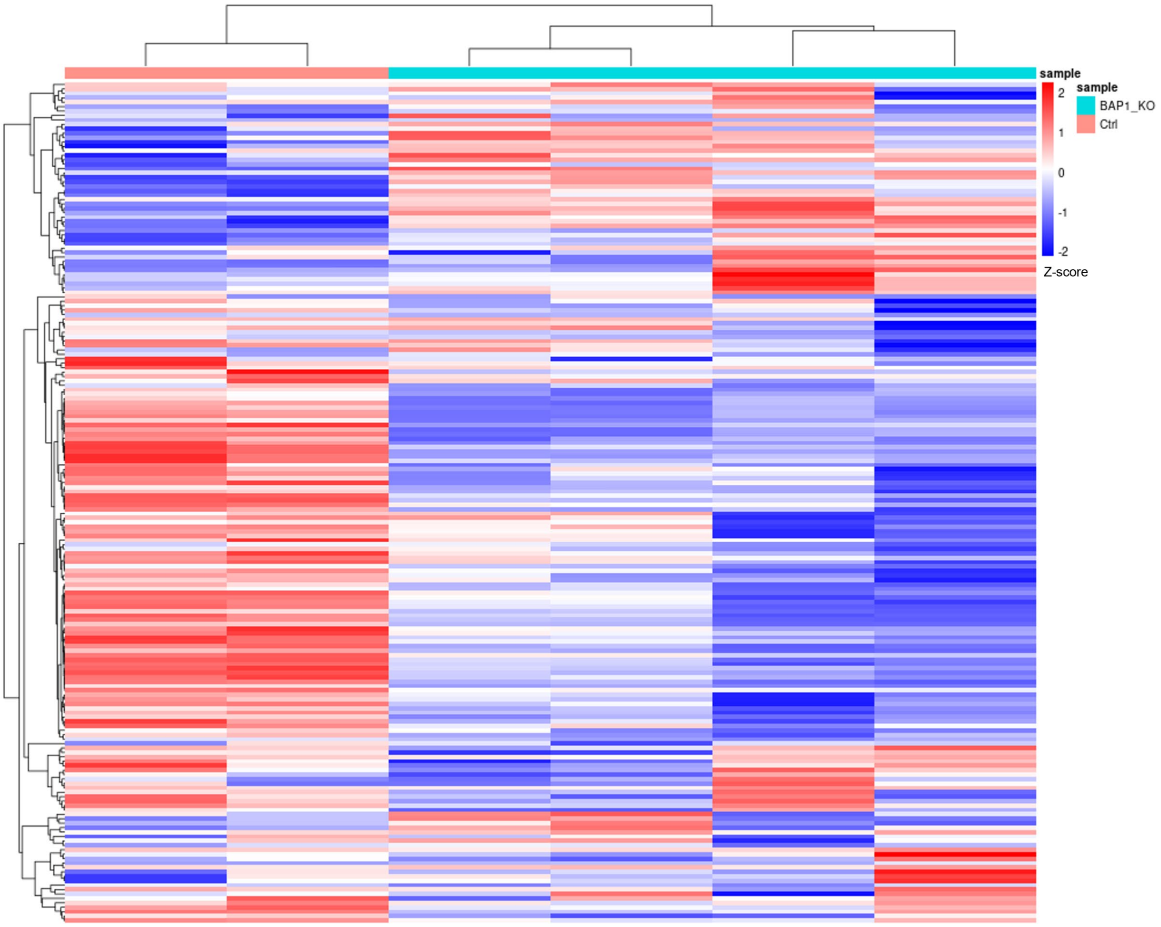


**Supplementary Figure S6**

**Supplementary Figure S6.** Heatmap with hierarchical clustering of mammosphere-cultured BAP1 KO and MCF10A-Cas9 cells based on RNA expression of glycosylation-associated genes (GO:0070085). Columns (samples) represent each replicate (n=2) of the MCF10A-Cas9 cells (Ctrl, pink) and BAP1 KO clones (BAP1_KO, blue), and rows represent the genes from the glycosylation pathway. Expression values are row-specific z-scores calculated from log10-transformed normalized counts from RNA-seq. BAP1 KOs exhibit clear differences in the expression of glycosylation-associated genes when compared to MCF10A-Cas9, showing overall reduced expression of genes in this pathway.

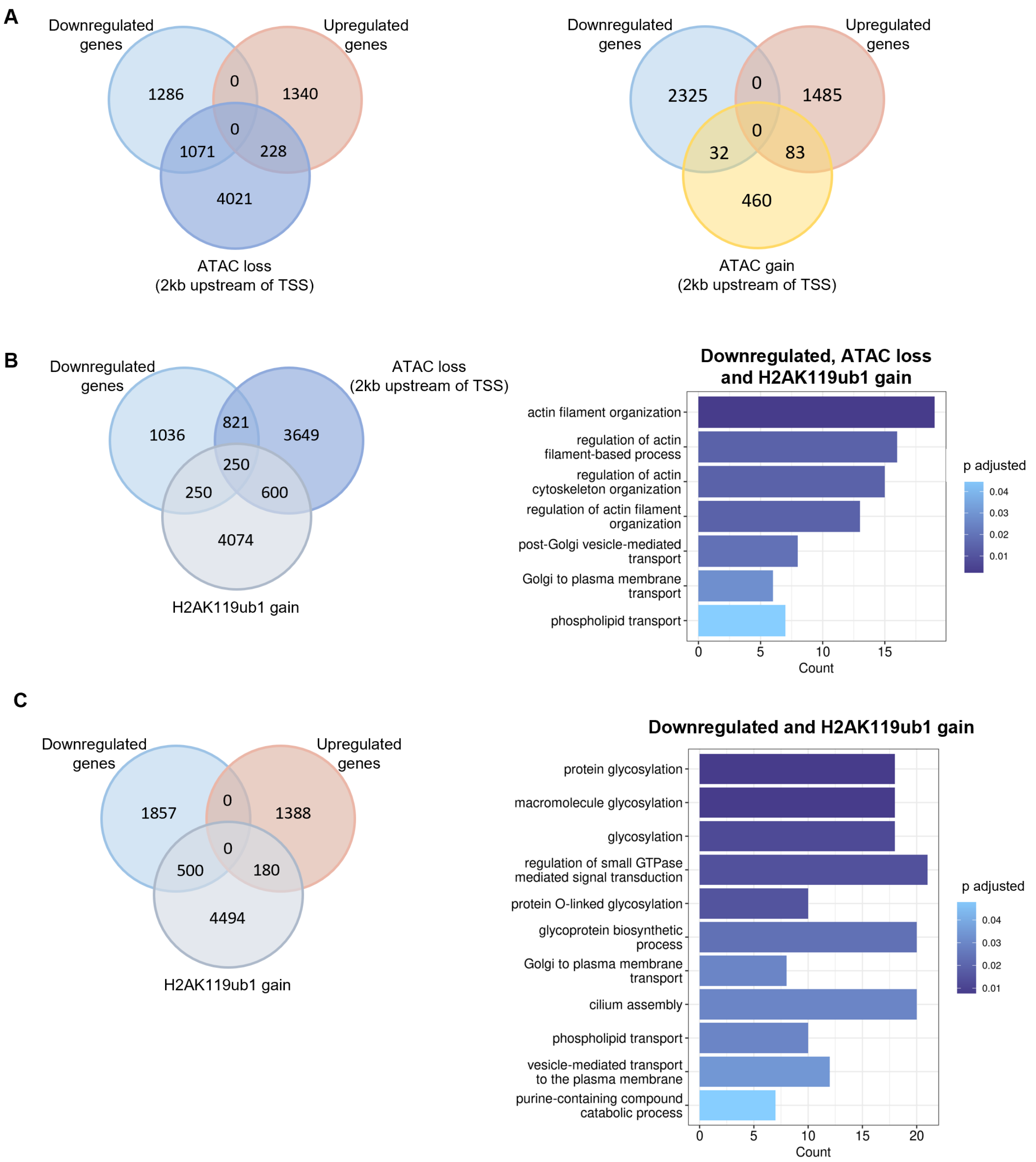


### Supplementary Figure S7

**Supplementary Figure S7.** (A) Venn diagram of genes associated with ATAC loss (FDR < 0.01, $\log_2FC < 0$, left) or ATAC gain (FDR < 0.01, $\log_2FC > 0$, right) within 2kb upstream of their TSSs and differential gene expression (FDR < 0.01) in mammosphere-cultured BAP1 KOs compared to MCF10A-Cas9. (B) Left: Venn diagram of genes associated with H2AK119ub1 gain (FDR < 0.05), ATAC loss (FDR < 0.01, $\log_2FC < 0$) within 2kb upstream of their TSSs and downregulation of gene expression (FDR < 0.01, $\log_2FC < 0$) in mammosphere-cultured BAP1 KOs compared to MCF10A-Cas9. Right: Gene ontology enrichment of 250 overlapping genes showing RNA downregulation, ATAC loss and H2AK119ub1 gain in mammosphere-cultured BAP1 KOs compared to MCF10A-Cas9 controls. All enriched ontologies are shown (p adjusted < 0.05). (C) Left: Venn diagram of overlap between genes associated with H2AK119ub1 gain (FDR <

0.05) and differential gene expression (FDR < 0.01) in BAP1 KOs compared to MCF10A-Cas9 control cells cultured as mammospheres. Right: Gene ontology enrichment of downregulated genes (FDR < 0.01, $log_2FC < 0$) showing H2AK119ub1 gain (FDR < 0.05) in BAP1 KOs compared to MCF10A-Cas9 control cells cultured as mammospheres. All enriched gene ontologies are shown (p adjusted < 0.05).

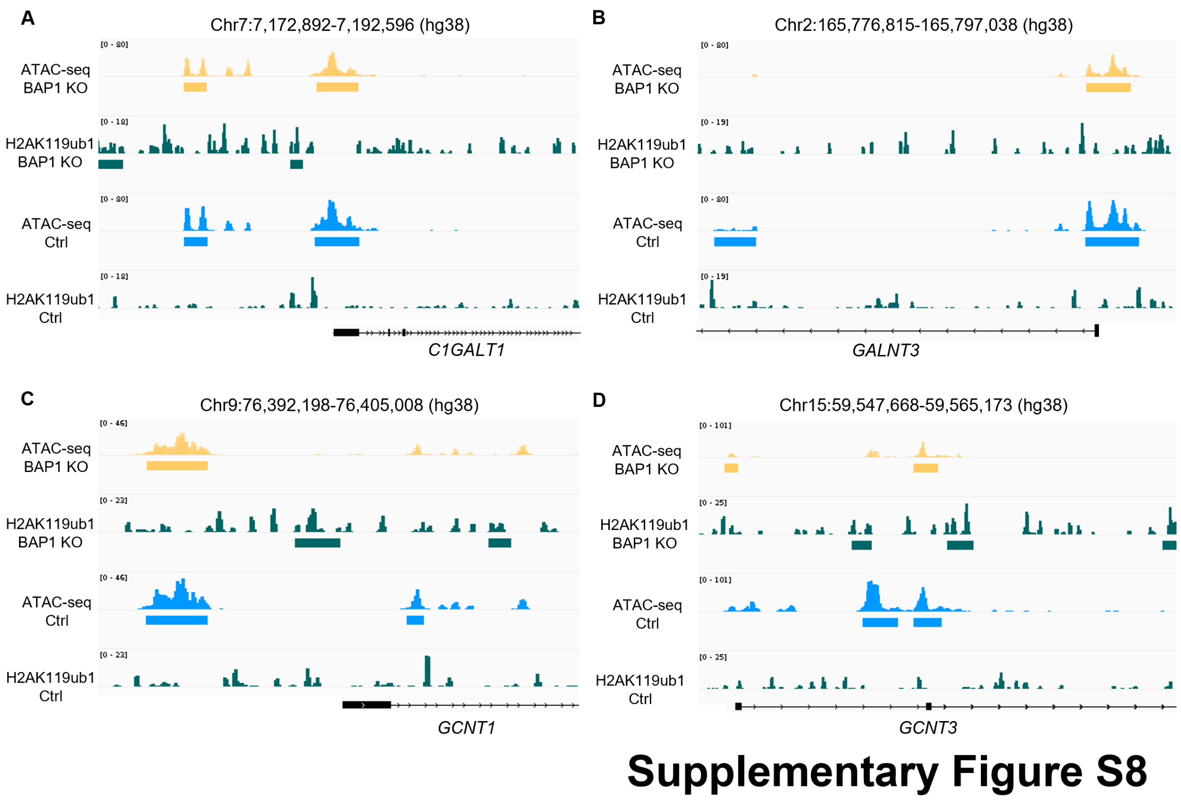


**Supplementary Figure S8.** Genome browser snapshot of ATAC-seq and H2AK119ub1 peaks in BAP1 KOs and MCF10A-Cas9 controls cultured as mammospheres, at the (A) *C1GALT1*, (B) *GALNT3*, (C) *GCNT1* and (D) *GCNT3* genes. Colored horizontal bars represent differential peaks identified in ATAC-seq (FDR < 0.01) and/or ChIP-seq (FDR < 0.05).

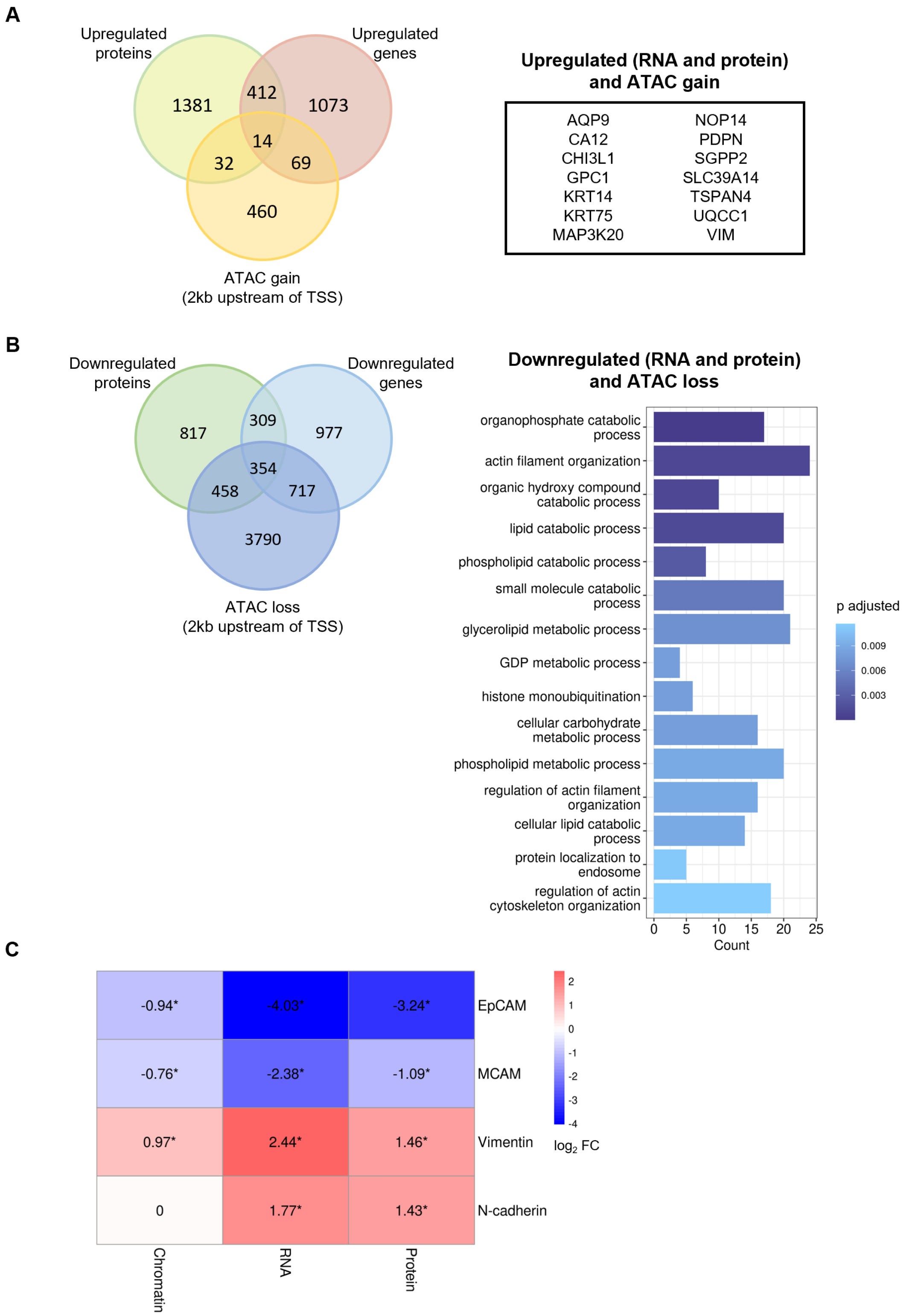


### Supplementary Figure S9

**Supplementary Figure S9.** (A) Left: Venn diagram of overlap between genes associated with ATAC gain (FDR < 0.01) within 2kb upstream of their TSSs, upregulated genes and proteins (log2FC > 0, FDR < 0.01) in BAP1 KOs compared to MCF10A-Cas9 control cells cultured as mammospheres. Right: List of the 14 upregulated genes showing ATAC gain and upregulation at the protein level in mammosphere-cultured BAP1 KOs compared to MCF10A-Cas9 controls.

(B) Left: Venn diagram of overlap between genes associated with ATAC loss (FDR < 0.01) within 2kb upstream of their TSSs, downregulated genes and proteins (log2FC < 0, FDR < 0.01) in BAP1 KOs compared to MCF10A-Cas9 control cells cultured as mammospheres. Right: Gene ontology enrichment of 354 downregulated genes showing ATAC loss and downregulation at the protein level in mammosphere-cultured BAP1 KOs compared to MCF10A-Cas9 controls. Top 15 gene ontologies are shown (p adjusted < 0.05). (C) Heatmap of EMT-associated genes showing consistent alteration in chromatin accessibility (ATAC-seq, 2kb upstream of TSSs, left), RNA expression (RNA-seq, middle) and protein levels (proteomics, right) in BAP1 KO mammospheres compared to MCF10A-Cas9 controls. Colors and numbers on the heatmap represent the log2FC between BAP1 KOs and controls for each omics analysis (red: ATAC gain or upregulation, blue: ATAC loss or downregulation, white: no change). Asterisks indicate that the change is significant (* FDR < 0.01). Where more than one ATAC-seq peak was annotated to a specific gene, only the closest peak to its TSS is shown in the heatmap. For the N-cadherin gene, the lack of ATAC-seq peaks identified within 2kb upstream of its TSS is represented by the white color and the number 0.

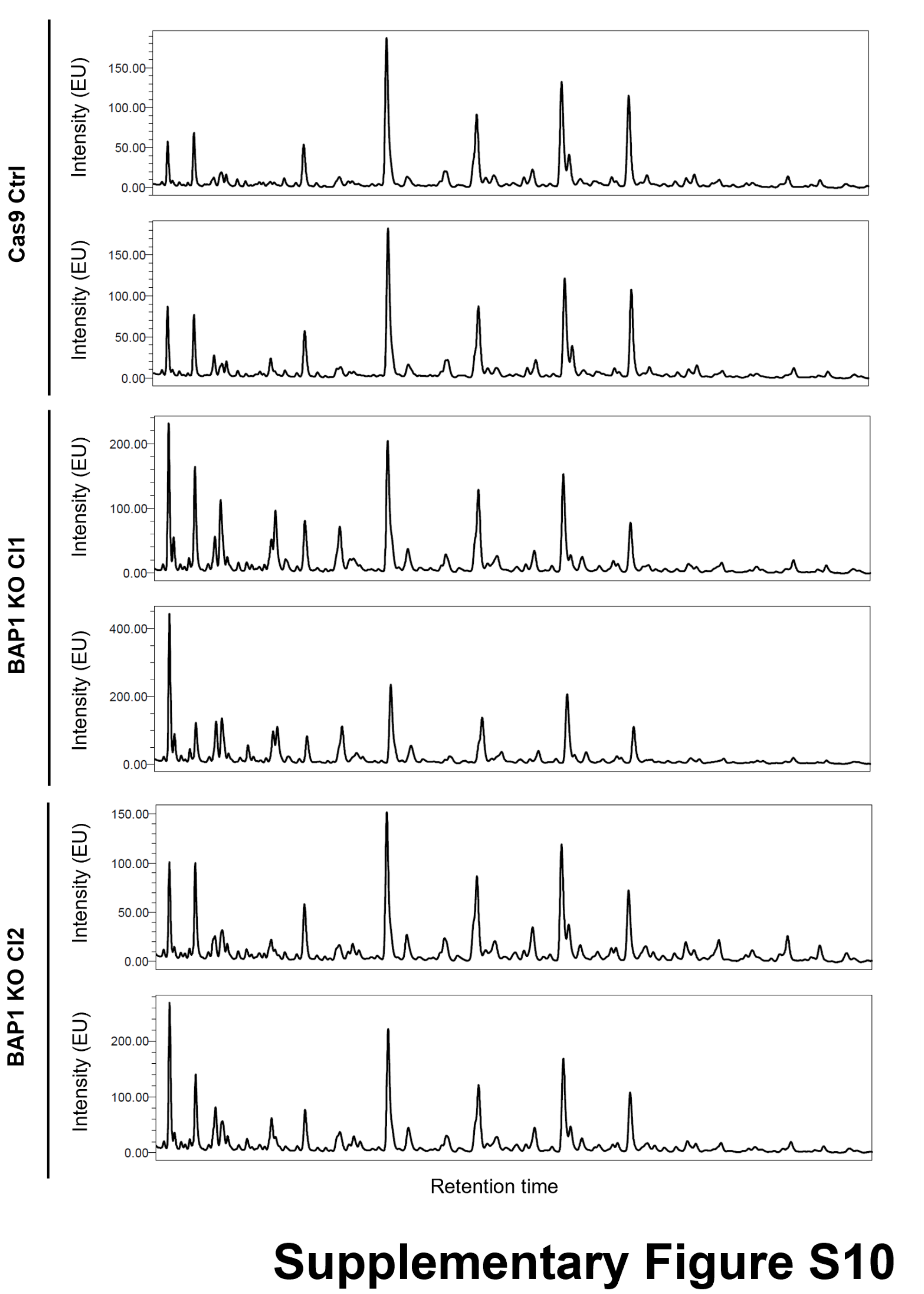


**Supplementary Figure S10.** Representative images of N-glycan profile chromatograms for MCF10A-Cas9 control and BAP1 KO cells cultured as mammospheres. Images shown are from two replicates out of the four analyzed in this assay. Peaks at earlier retention times represent less complex N-glycans, while peaks at later retention times represent more complex N-glycan structures.

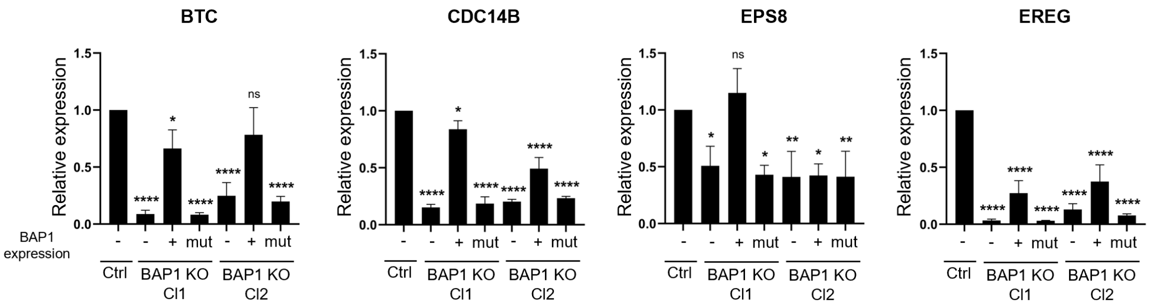


### Supplementary Figure S11

**Supplementary Figure S11.** Expression of cell cycle-associated genes *BTC*, *CDC14B*, *EPS8* and *EREG* in MCF10A-Cas9 (Ctrl), BAP1 KO cells and BAP1 rescues (wildtype, +; and mutant, mut) cultured as mammospheres. Significance analysis performed by Dunnett's test compared to control (Ctrl) sample (n=3, * $P < 0.05$, ** $P < 0.01$, **** $P < 0.0001$, ns: not significant).

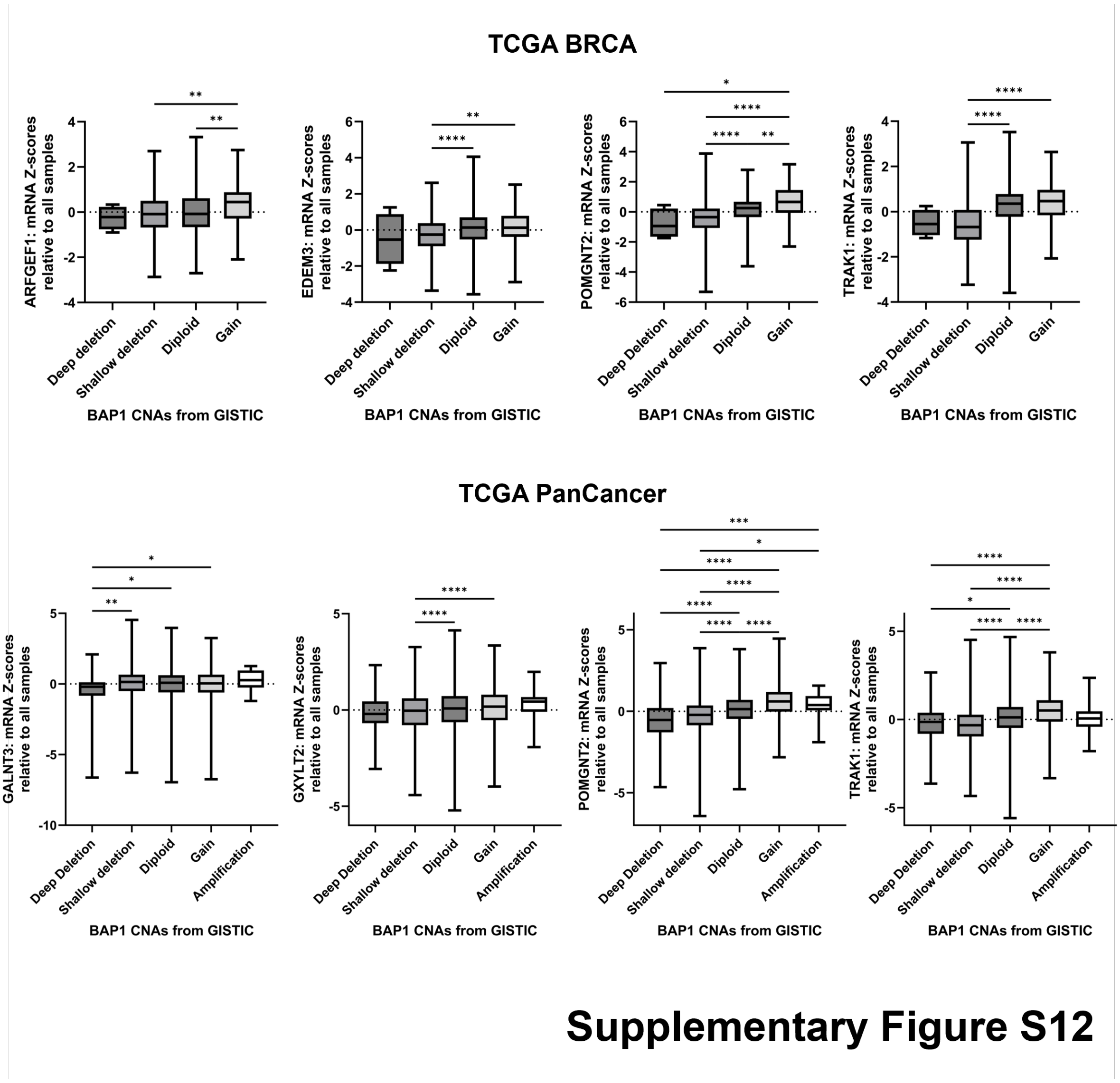


**Supplementary Figure S12.** Boxplots for correlation between BAP1 copy number alterations (CNAs) and mRNA expression of glycosylation-associated genes in breast cancer tumors (BRCA, top) and tumors from 32 cancer types (PanCancer, bottom), using data from TCGA (The Cancer Genome Atlas). mRNA expression levels are shown as Z-scores relative to all samples. BAP1 copy numbers are derived from the GISTIC algorithm, and are distributed as follows. TCGA BRCA (total 1,068 samples): deep deletion (n=4), shallow deletion (n=339), diploid (n=640), gain (n=85). TCGA PanCancer (total 9,889 samples): deep deletion (n=77), shallow deletion (n=3,152), diploid (n=5,820), gain (n=820), amplification (n=20). Plots and statistical analysis were generated in GraphPad Prism v10.3.1 with TCGA data obtained from cBioPortal (https://www.cbioportal.org/). Significance analysis performed by Kruskal-Wallis test with Dunn's multiple comparisons (* $P < 0.05$, ** $P < 0.01$, *** $P < 0.001$, **** $P < 0.0001$).

#### SUPPLEMENTARY TABLES

**Supplementary Table S1. gRNA sequences used for targeting of the *BAP1* gene through a CRISPR/Cas9 loss-of-function approach.**

| Gene name | NCBI Gene ID | gRNA ID | gRNA sequence | Chromosome | Exon |
|---|---|---|---|---|---|
| BAP1 | 8314 | 1 | TCAAATGGATCGAAGAGCGC | chr3 | Exon 4 |
| BAP1 | 8314 | 2 | CACGGACGTATCATCCACCA | chr3 | Exon 4 |
| BAP1 | 8314 | 3 | CGACCTTCAGAGCAAATGTC | chr3 | Exon 3 |
| BAP1 | 8314 | 4 | CGCATGAAGGACTTCACCAA | chr3 | Exon 5 |

**Supplementary Table S2. Primer sequences used in RT-qPCR analyses.**

| Target gene | Forward primer sequence (5'-3') | Reverse primer sequence (5'-3') | Reference |
|---|---|---|---|
| BTC | CCTCTTCGGAAACGTCGTAAAAG | CTTGCCACCAACCTGGAGGTAA | - |
| C1GALT1 | TCATCCCTTTGTGCCAGAAC | CAAGATCAGAGCAGCAACCA | - |
| CDC14B | AGGATGTATGATGCCAAACGC | GCTGCTGTCATCCTGTAATGC | PMID: 21379580 |
| CDH1 (E-cadherin) | TCCTGGGCAGAGTGAATTTTG | CTGTAATCACACCATCTGTGC | - |
| CDH2 (N-cadherin) | CCACAATCCTGTCCACATCT | TTCGGGTAATCCTCCCAAAT | - |
| EPCAM | AGAACCTACTGGATCATCATTGAACTAA | CGCGTTGTGATCTCCTTCTG | - |
| EPS8 | GATGGAGGAAGTGCAAGATG | GACTGTAACCACGTCTTCACA | PMID: 29192326 |
| EREG | CTTATCACAGTCGTCGGTTCCAC | GCCATTCAGACTTGCGGCAACT | - |
| GALNT3 | CTCTATGTCTGGATGTTGG | TCATGTTGAGCAGAGTATTC | PMID: 18976705 |
| GAPDH | GTCTCCTCTGACTTCAACAGCG | ACCACCCTGTTGCTGTAGCCAA | - |
| GCNT1 | CTGGAAACGGAGAGGATGCCAT | CACGAAGTAGGCACTGCCAGAA | - |
| ST6GALNAC1 | CGAAATAGGAGGCCTTCAGA | AGAGAGTGAGGTTGGGCAGA | PMID: 30996686 |
| VIM (Vimentin) | AAAGTGTGGCTGCCAAGAAC | AGCCTCAGAGAGGTCAGCAA | - |
| ZEB1 | GCTGGGAGGATGACAGAAAG | TGCATCTGACTCGCATTCAT | - |
| ZEB2 | GGCTCCATGTTCATAGCATAGT | CCTCCTCTTGTCATCTGTACTTTC | - |

**Supplementary Table S3. Primer sequences used in pyrosequencing analyses.**

| Target gene | Forward primer sequence (5'-3') | Reverse biotinylated primer sequence (5'-3') | Sequencing primer (5'-3') |
|---|---|---|---|
| CDH1 | TTAGTTTYGTTTTGGGGAGGGG | CCCCCACTCCCATCACTAAAA | GTTAGTTTAGATTTTAGTT |
| ZEB1 | GATTTTTAAGAGAGGGGTAATAAA | CCACACAAAATCTAAAATTACCA | TTTTTTTTTTTGGGATG |

**Supplementary Table S4. Enriched gRNAs in the cancer stem cell-like population of MCF10A cells infected with the epigenetic regulator gene (ERG) library compared to the bulk of cells on the day of sorting.**

| Gene | Enrichment rank | Number of enriched gRNAs | p-value |
|---|---|---|---|
| KDM6A | 1 | 4 | 0.005 |
| ASXL2 | 2 | 4 | 0.0074 |
| BAP1 | 3 | 3 | 0.0421 |
| PRDM9 | 4 | 2 | 0.0421 |
| CBX4 | 5 | 4 | 0.0421 |
| KAT6B | 6 | 3 | 0.0421 |

**Supplementary Table S5. Frequently mutated epigenetic regulator genes across BCs and exclusive to specific BC subtypes.** Abbreviations: ChRC: chromatin remodeling factors; DM: DNA methylation; H: histone; HA: histone acetylation; HM: histone methylation; w: writer; e: eraser; r: reader; pSNA: proportion of single nucleotide alterations; ERpHER2n: luminal A; ERpHER2p: luminal HER; ERnHER2p: HER2-enriched; TNBC: triple negative breast cancer. Epigenetic regulator genes identified as exclusively mutated in one subtype (pSNA cutoff > 0.01) are highlighted in red.

Table supplied as Excel file.

**Supplementary Table S6. Genomic compartment distribution of differential ATAC-seq peaks (FDR < 0.01) identified in BAP1 KOs compared to MCF10A-Cas9 controls cultured as mammospheres.**

| | ATAC gain | | ATAC loss | |
|---|---|---|---|---|
| **Genomic compartment** | **Number of peaks** | **Percentage** | **Number of peaks** | **Percentage** |
| Upstream 10kb | 402 | 4.94 | 1440 | 5.08 |
| Upstream 2kb | 343 | 4.22 | 1385 | 4.89 |
| Promoter | 253 | 3.11 | 4455 | 15.72 |
| Gene body | 4059 | 49.91 | 13055 | 46.06 |
| Exon | 461 | 5.67 | 1880 | 6.63 |
| Intergenic | 2615 | 32.15 | 6130 | 21.63 |

**Supplementary Table S7. Chromatin state distribution of differential ATAC-seq peaks (FDR < 0.01) identified in BAP1 KOs compared to MCF10A-Cas9 controls cultured as mammospheres.**

| | ATAC gain | | ATAC loss | |
|---|---|---|---|---|
| **Chromatin state** | **Number of peaks** | **Percentage** | **Number of peaks** | **Percentage** |
| Active TSS | 45 | 0.56 | 3699 | 13.11 |
| Flanking TSS | 287 | 3.55 | 1723 | 6.11 |
| Flanking transcription | 2 | 0.03 | 28 | 0.1 |
| Strong transcription | 833 | 10.32 | 3402 | 12.06 |
| Weak transcription | 129 | 1.6 | 545 | 1.93 |
| Genic enhancer | 59 | 0.73 | 396 | 1.4 |
| Enhancer | 2743 | 33.97 | 10376 | 36.78 |
| ZNF repeats | 9 | 0.11 | 29 | 0.1 |
| Heterochromatin | 337 | 4.17 | 458 | 1.62 |
| Bivalent/poised TSS | 4 | 0.05 | 52 | 0.18 |
| Flanking bivalent TSS | 16 | 0.2 | 52 | 0.18 |
| Bivalent enhancer | 23 | 0.29 | 52 | 0.18 |
| Repressed Polycomb | 27 | 0.33 | 42 | 0.15 |
| Weak repressed Polycomb | 577 | 7.15 | 1260 | 4.47 |
| Quiescent regions | 2985 | 36.96 | 6100 | 21.62 |
| Non-categorized (NA) | 57 | - | 131 | - |

**Supplementary Table S8. Transcription factor motif enrichment analysis in differentially accessible chromatin regions (FDR < 0.01) within 2kb upstream of TSSs in BAP1 KOs compared to MCF10A-Cas9 controls cultured as mammospheres.**

Table supplied as Excel file.
